# Parkinson’s VPS35[D620N] mutation induces LRRK2 mediated lysosomal association of RILPL1 and TMEM55B

**DOI:** 10.1101/2023.06.07.544051

**Authors:** Prosenjit Pal, Matthew Taylor, Pui Yiu Lam, Francesca Tonelli, Chloe A. Hecht, Pawel Lis, Raja S. Nirujogi, Toan K. Phung, Emily A. Dickie, Melanie Wightman, Thomas Macartney, Suzanne R. Pfeffer, Dario R. Alessi

**Affiliations:** MRC Protein Phosphorylation and Ubiquitylation Unit, University of Dundee; Department of Biochemistry, Stanford University School of Medicine, Stanford, CA; Aligning Science Across Parkinson’s (ASAP) Collaborative Research Network, Chevy Chase, MD, 20815

**Author notes:** Corresponding author. (P.P.); (D.R.A.). These authors contributed equally to this work (Prosenjit Pal, Matthew Taylor, Pui Yiu Lam).

## Abstract

The Parkinson’s VPS35[D620N] mutation causes lysosome dysfunction enhancing LRRK2 kinase activity. We find the VPS35[D620N] mutation alters expression of ∼350 lysosomal proteins and stimulates LRRK2 phosphorylation of Rab proteins at the lysosome. This recruits the phosphoRab effector protein RILPL1 to the lysosome where it binds to the lysosomal integral membrane protein TMEM55B. We identify highly conserved regions of RILPL1 and TMEM55B that interact and design mutations that block binding. In mouse fibroblasts, brain, and lung, we demonstrate that the VPS35 [D620N] mutation reduces RILPL1 levels, in a manner reversed by LRRK2 inhibition. Knock-out of RILPL1 enhances phosphorylation of Rab substrates and knock-out of TMEM55B increases RILPL1 levels. The lysosomotropic agent LLOMe, also induced LRRK2 kinase mediated association of RILPL1 to the lysosome, but to a lower extent than the D620N mutation. Our study uncovers a pathway through which dysfunctional lysosomes resulting from the VPS35[D620N] mutation recruit and activate LRRK2 on the lysosomal surface, driving assembly of the RILPL1-TMEM55B complex.

## Introduction

Mutations that increase the kinase activity of the leucine-rich repeat kinase-2 (LRRK2) represent one of the most common inherited causes of Parkinson’s disease (*1, 2*) and have also been linked to inflammatory bowel disease (*3, 4*). LRRK2 is a large 2527 residue multidomain protein consisting of two catalytic domains: a Roco type GTPase in addition to a protein kinase (*5*). LRRK2 phosphorylates a subgroup of Rab GTPase proteins (Rab1, Rab3, Rab8, Rab10, Rab12, Rab29, Rab35, and Rab43) (*6, 7*), that coordinate membrane homeostasis and endocytic and exocytic pathways (*8*). LRRK2 phosphorylates Rab substrates at a conserved Ser or Thr site lying at the center of the effector-binding switch-II motif (Thr72 for Rab8A, Thr73 for Rab10, and Ser106 for human Rab12) (*6, 7*). This reaction is counteracted by the PPM1H phosphatase that efficiently dephosphorylates Rab proteins (*9, 10*). LRRK2 phosphorylation of Rab proteins impacts their ability to interact with its cognate effectors, for example, phosphorylation of Rab8 blocks interactions with Rabin-8 (a GDP/GTP exchange factor) and GDI (GDP dissociation inhibitor) that shuttles Rab proteins between membranes (*6, 7*).

Four scaffolding proteins, namely RILPL1, RILPL2, JIP3 and JIP4, interact specifically with LRRK2 phosphorylated Rab8A and Rab10 with higher affinity than dephosphorylated Rab proteins (*6, 11, 12*). This interaction is mediated by an α-helical RH2 motif possessing conserved basic residues that form ionic interactions with the LRRK2 phosphorylated Switch-II motif residue (*11*). The interaction of RILPL1 with phosphorylated Rab proteins interferes with ciliogenesis (*6, 13*) and in cholinergic neurons in the striatum leads to disruption of a Sonic hedgehog neuro-protective circuit that supports dopaminergic neurons, providing a pathway by which LRRK2 may be linked to Parkinson’s disease pathology (*14*).

Rab proteins also play a key role in controlling LRRK2 kinase activity by binding to the N-terminal ARM domain and recruiting LRRK2 to membranes where it becomes activated (*15, 16*). Recent work has pinpointed 3 Rab binding sites within the LRRK2 ARM domain. Site-1 binds to dephosphorylated Rab29, Rab32 as well as Rab8 and possibly Rab10 (*17-19*). Site-2 interacts specifically to LRRK2-phosphorylated Rab8 and Rab10 in a feed-forward pathway that drives membrane recruitment and activation of LRRK2 (*18*). Site-3 interacts with Rab12 (*20*) and ablation of this site or knock-out of Rab12 has the largest effect in regulating the basal activity of LRRK2 as judged by its ability to phosphorylate Rab10 (*20, 21*).

Lysosomal dysfunction is strongly associated with PD (*22, 23*). LRRK2 and other PD-associated genes including GBA1 (*24*), ATP13A2 (*25*) and TMEM175 (*26*), play a critical role in controlling lysosome homeostasis and function. Elevated LRRK2 kinase activity reduces lysosomal degradative activity and autophagic flux in a manner that is counteracted by LRRK2 inhibitors (*27-29*). Furthermore, damage of lysosomes following infection (*30*) or treatment with agents such as LLOME induce recruitment of LRRK2 to lysosomal membranes (*12, 31*). At the lysosome LRRK2 is activated and found to phosphorylate Rab proteins thereby recruiting JIP4, which promotes formation of tubular structures that release membranous content from lysosomes (*12*). Recent work reveals that LRRK2 negatively regulates lysosomal degradative activity in macrophages and microglia via a transcriptional mechanism involving transcription factor E3 (TFE3) (*32*). Depletion of LRRK2 and inhibition of LRRK2 kinase activity both enhance lysosomal proteolytic activity and increase the expression of multiple lysosomal hydrolases (*32*). Other work has revealed that LRRK2 kinase activity controls PD relevant lipids such as bis(monoacylglycerol)phosphates as well as glycosphingolipids at the lysosome (*33, 34*).

The VPS35 component of the retromer complex transports select endosomal cargo proteins between endosomal compartments and the Golgi and has been linked to Parkinson’s (*35, 36*) as well as Alzheimer’s diseases (*37*). The D620N mutation in VPS35 causes autosomal dominant Parkinson’s and stimulates the LRRK2 pathway via an unknown mechanism (*38*). VPS35[D620N] knock-in cells and tissues display markedly enhanced Rab phosphorylation to a higher level than observed with LRRK2 PD pathogenic mutations (*38, 39*) and this mutation has been proposed to lead to lysosomal dysfunction (*40, 41*). It is possible that D620N VPS35 disruption of selective endosomal cargo trafficking triggers lysosomal dysfunction, thereby activating LRRK2.

In this study we sought to investigate the impact that elevated LRRK2 signalling has on lysosomal protein content employing LysoTag immunoprecipitation coupled to mass spectrometry. We have uncovered a new pathway by which lysosomal stress or dysfunction resulting from the VPS35[D620N] mutation or the lysosomotropic agent LLOMe induces lysosomal recruitment of LRRK2, resulting in phosphorylation of Rab proteins which triggers recruitment of RILPL1. Our data suggest that RILPL1 then interacts via its conserved C-terminal region with a conserved domain of TMEM55B, an integral lysosomal membrane protein. Our data provide new insights into a pathway by which LRRK2 communicates with the lysosome.

## Results

### Impact of VPS35[D620N] mutation on lysosomal protein content

To study the impact that the VPS35[D620N] mutation has on the lysosome, we first performed a LysoTag immunoprecipitation in lysates from littermate matched wild type and homozygous knock-in VPS35[D620N] mouse embryonic fibroblasts (MEFs) transduced to stably express LysoTag (TMEM192-3xHA) (*42*), employing a workflow described in Figure 1A. Immunoblot analysis confirmed previous data that the VPS35[D620N] mutation enhanced LRRK2-mediated Rab10 phosphorylation ∼4-fold (Fig. 1B). Data-independent acquisition (DIA) mass spectrometry (MS) with equal protein amounts (4μg) of whole cell lysates (Fig. 1C) and isolated lysosomes was undertaken, with 6 replicates (Fig. 1D). MS data was searched through DIA-NN (*43*), and visualized using a new interactive visualization tool called Curtain (https://curtain.proteo.info) in which the data can be analyzed using the web links provided in the Figure legend. The experiments revealed that in whole cell lysates, the D620N mutation altered expression >2-fold of 363 proteins, with 70 increasing (0.82%) and 293 decreasing (3.44%) (Fig. 1C and Table S1). Similarly, for the LysoTag immunoprecipitation, 81 proteins increased and 136 decreased >2-fold (Fig. 1D; fig. S1, A to C and Table S1). Violin plots for the top 12 proteins whose levels increase with the D620N mutation (including Cathepsin C, Cathepsin K, SLC38A4 and GJB2) and the top 12 that decrease (including THY1, HSPA1A/B, MT2 and SFRP2) are shown in Figure S2A, B. We undertook gene ontology metascape analysis (*44*) of the proteins that were increased or decreased in VPS35[D620N] compared to wild type MEFs in whole lysates (fig. S3A and Table S1) and LysoTag immunoprecipitates (fig. S3B and Table S1). In whole cell lysates, proteins that changed in VPS35[D620N] MEFs impacted a wide range of biology including response to virus, cell proliferation, adhesion, nucleosome and extracellular matrix. The top proteins changing in the VPS35[D620N] lysosomes immunoprecipitates included those impacting extracellular matrix, glycosaminoglycan binding, vacuolar membrane, transmembrane transport and anion channel activity.

**Fig 1.**
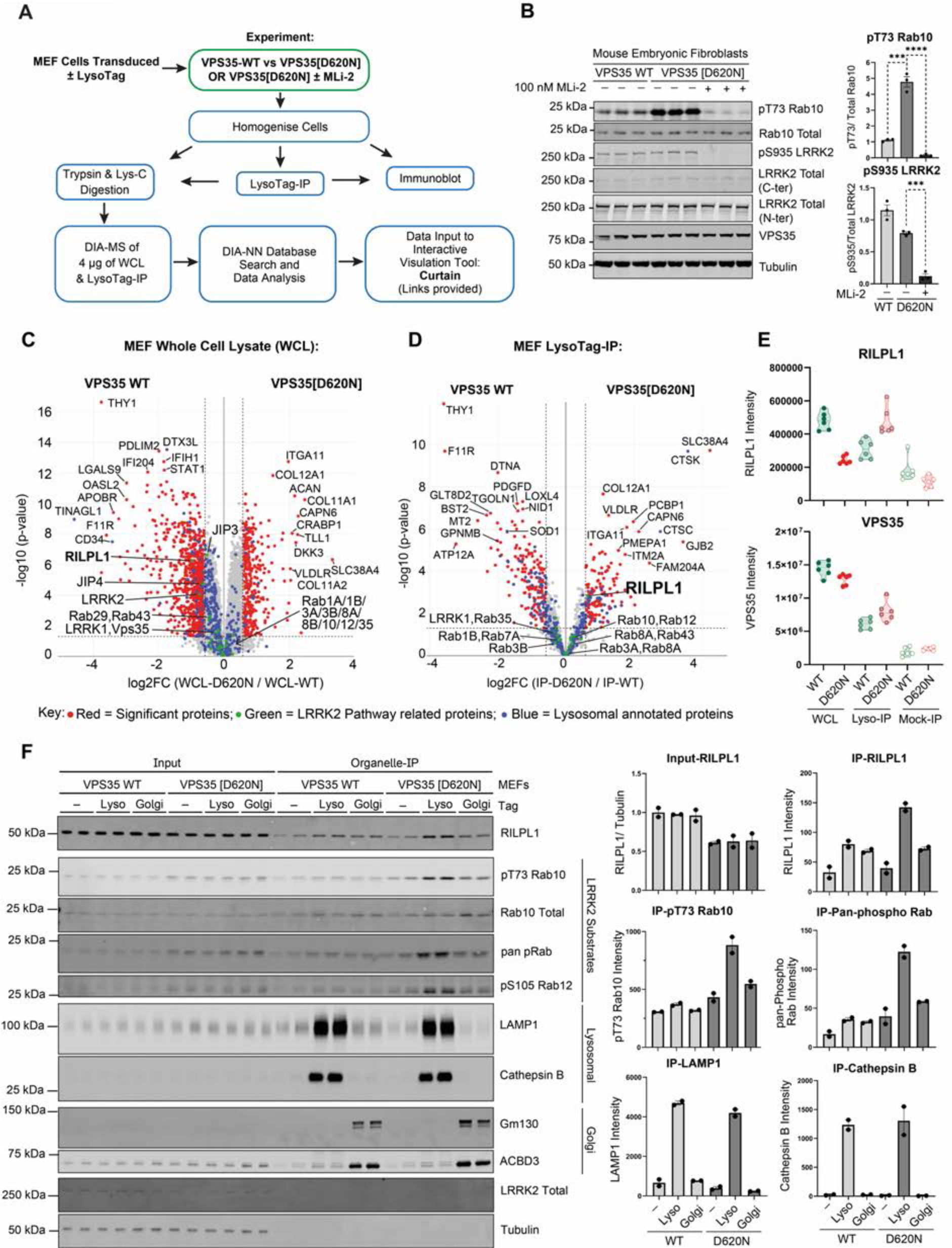
Phospho-Rab and RILPL1 enrichment at VPS35[D620N] mutant lysosome. (A) Schematic workflow of the LysoTag-IP methodology in MEFs used for both immunoblotting and MS analysis. (B) Littermate-matched wild type (WT) and VPS35[D620N] homozygous knock-in (KI) MEFs were treated ± 100 nM MLi-2 for 2 h prior to lysis. Lysate were subjected to quantitative immunoblot analysis using the LI-COR Odyssey CLx Western Blot imaging and the indicated antibodies. Technical replicates represent cell extract obtained from different dishes of cells. Quantitation of immunoblotting data (performed using ImageStudioLite software version 5.2.5, RRID:SCR_013715; http://www.licor.com/bio/products/software/image_studio_lite/) is shown as mean ± SEM. Data were analyzed using two-tailed unpaired t-test (*** p< 0.001, **** p< 0.001). (C-D) Littermate WT and VPS35[D620N] homozygous KI MEFs were transduced ± LysoTag (TMEM192-3xHA) and subjected to homogenization. A sample of the homogenate was removed and designated whole cell lysate (WCL). The remaining homogenate was subjected to anti-HA LysoTag-IP. Experiment was performed in 6 technical replicates. DIA-MS experiment was performed on an Orbitrap Exploris 480 MS. The volcano plots show the proteome changes of MEFs VPS35-WT versus VPS35[D620N] in (C) whole cell lysate (Curtain link: https://curtain.proteo.info/#/0e673d58-d8f2-4368-996b-0869d5513d46) and (D) lysosome (Curtain link:https://curtain.proteo.info/#/0e673d58-d8f2-4368-996b-0869d5513d46) respectively (Table S1). In (D), only proteins with fold change > 1.5 compared to an Mock-IP using cells expressing no LysoTag (fig. S1A) were highlighted in the Volcano plot. The red dots represent the significant differentiated proteins with fold change > 1.5 fold and p-value < 0.05; the green dots represent the LRRK2 pathway related proteins; and the blue dots represent the lysosomal annotated proteins. (E) Violin plots of RILPL1 and VPS35 levels derived from experiment in (C, D). (F) Littermate-matched WT and VPS35[D620N] homozygous knock-in MEFs transduced ± either the LysoTag (TMEM192-3xHA) or GolgiTag (TMEM115-3xHA) were subjected to homogenization and organelle isolation as in (C-D). 2 μg of both the immunoprecipitate and respective input (WCL) was subjected to immunoblot analysis using indicated antibodies as in (B). Each lane indicates a sample derived from a different dish of cells. Quantification of immunoblotting is shown as mean ± SEM.

### LRRK2 activity drives the recruitment of RILPL1 to lysosomes in VPS35[D620N] cells

We searched the data for known LRRK2 pathway components and found one protein, the phospho-Rab effector protein RILPL1 (*6, 11*), whose levels were enriched in VPS35[D620N] compared to wild type lysosomes (Fig. 1E). To investigate this further, we performed LysoTag (*42*) and GolgiTag (*45*) immunoprecipitations in parallel from littermate-matched wild type and homozygous VPS35[D620N] knock-in MEFs (Fig. 1F). This revealed that the VPS35[D620N] mutation enhanced recruitment of RILPL1 specifically to the lysosome, but not the Golgi (Fig. 1F). We also observed increased phosphorylation of Rab10 (Thr73) and Rab12 (Ser105), as well as phosphorylated Rab proteins detected using a pan phospho-specific Rab antibody, at the VPS35[D620N] lysosome, but not at the Golgi (Fig. 1F). Note that recruitment of LRRK2 to the lysosome by the VPS35[D620N] mutation was not detected by immunoblotting experiments, consistent with the mass spectrometry data shown in Figure 1D. It is possible that LRRK2 dissociates from lysosomes following cell lysis and the wash steps in the LysoTag immunoprecipitation.

We next undertook LysoTag immunoprecipitations from VPS35[D620N] knock-in MEFs treated for 48h ± 100 nM MLi-2, a highly specific and well characterized LRRK2 inhibitor (*46*). Immunoblotting revealed that MLi-2 reduced levels of RILPL1 at the VPS35[D620N] lysosome to background levels observed in the wild-type immunoprecipitate (Fig. 2A). As expected, MLi-2 also ablated Rab10 phosphorylation in whole cell extract as well as the lysosome (Fig. 2A). MS analysis revealed that MLi-2 did not significantly alter the expression of any protein in the whole cell lysate >2-fold (Fig. 2B and Table S2). For the LysoTag immunoprecipitation, RILPL1 was the clearcut protein whose association with the lysosome was most reduced (∼2.5-fold) by MLi-2 treatment (Fig. 2C, fig. S4 and Table S2). Rab43, a LRRK2 substrate, was also reduced, although at borderline statistical significance (Fig. 2C). To our knowledge neither RILPL1 nor Rab43 have previously been associated with the lysosome. The lysosomal levels of several other proteins including LRRK2, ATP6V0D1, Laptm4a and VPS28 were moderately increased (∼1.5-fold) following MLi-2 treatment (Fig. 2C and fig. S4). We also investigated the levels of the top 24 proteins analyzed in Figure S2 whose expression was most impacted by the VPS35[D620N] mutation in the LysoTag-IP. We found that MLi-2 had little effect on most proteins, however the MT2 membrane-anchored serine protease, whose lysosomal levels were reduced in the VPS35[D620N] lysosomes, were moderately increased following MLi-2 treatment (fig. S5, A and B;Table S2).

**Fig 2.**
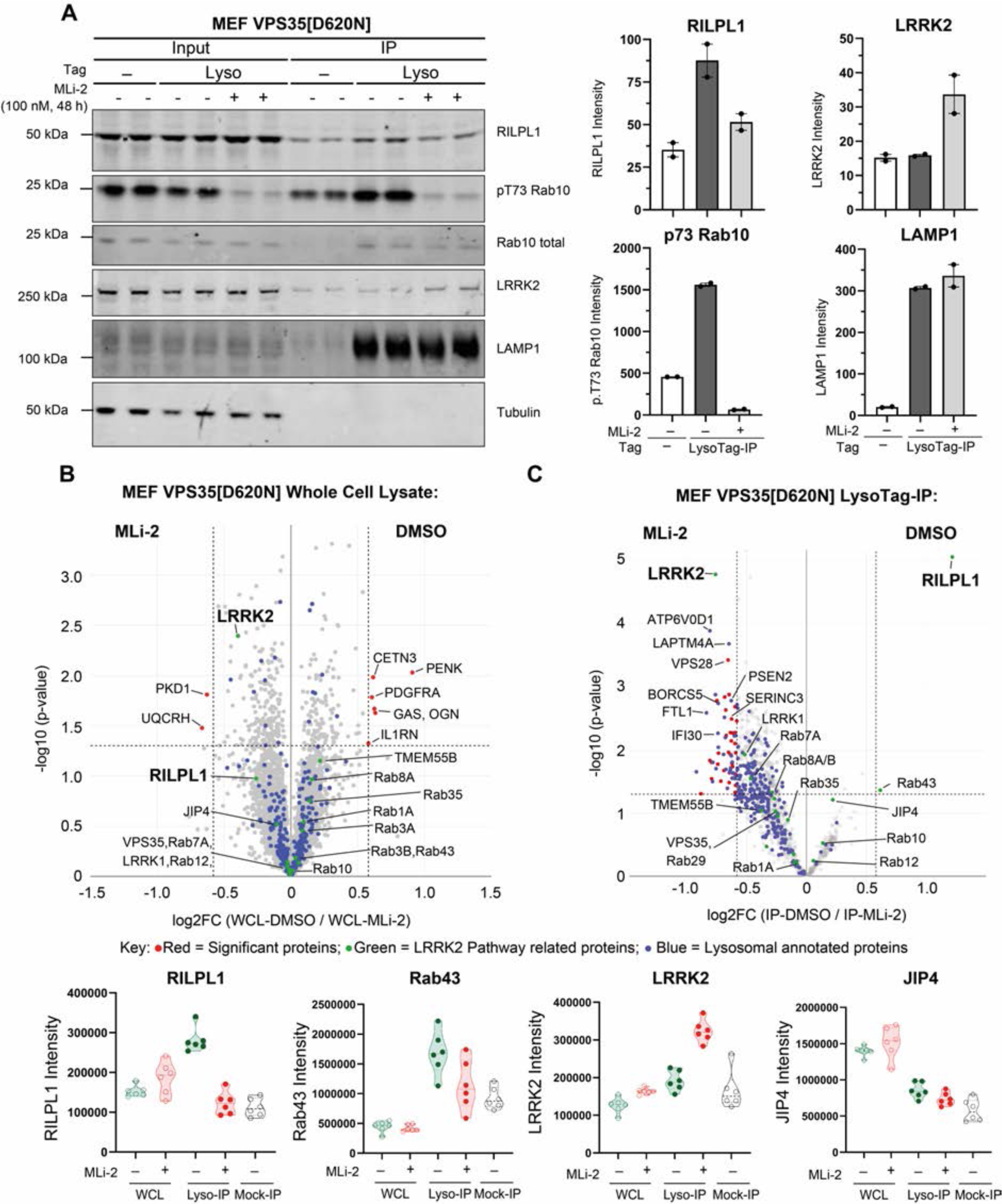
Lysosomal recruitment of RILPL1 is dependent on LRRK2 kinase activity. (A) VPS35[D620N] homozygous knock-in MEFs transduced ± LysoTag (TMEM192-3xHA) were subjected to ± 100 nM MLi-2 treatment for 48 h prior to homogenization. A sample of the homogenate was removed and labeled input and the remainder subjected to LysoTag-IP. 2 μg of both the input and immunoprecipitate was subjected to quantitative immunoblot analysis using the LI-COR Odyssey CLx Western Blot imaging system and indicated antibodies. Each lane represents a sample derived from a different dish of cells. Quantitation of immunoblotting data (performed using ImageStudioLite software version 5.2.5, RRID:SCR_013715; http://www.licor.com/bio/products/software/image_studio_lite/) is shown as mean ± SEM. (B-C) Immunoprecipitate and input (WCL) samples generated as in (A) were performed in 6 technical replicates and were subjected to DIA-MS analysis. The volcano plots show the proteome changes of MEFs D620N with DMSO versus MLi-2 treatment in whole cell lysates (Curtain link: https://curtain.proteo.info/#/d864df78-e2a5-4a64-99fb-8c5d8b2e1ab8) (B) and lysosomes (Curtain link: https://curtain.proteo.info/#/062e0a64-1fd3-4cc1-833a-a5b007d95a3c) (Table S2) (C). Proteins with fold change > 1.5 compared to Mock-IP samples are highlighted in the volcano plot (fig. S2A). The red dots represent the significant differentiated proteins with fold change > 1.5-fold and p-value < 0.05; the green dots represent the LRRK2 pathway related proteins; and the blue dots represent the lysosomal annotated proteins. (D) Violin plots of the levels of the indicated proteins.

We next investigated the levels of the set of lysosomal proteins (Cathepsin B, Cathepsin C, Cathepsin D, Cathepsin L, GBA, LAMP1, TFE3 and TFEB) that were previously reported to increase following inhibition or depletion of LRRK2 in macrophages and microglia (*32*). In wild type versus VPS35[D620N] and VPS35[D620N] ± MLi-2 MEF datasets, only cathepsin L was moderately decreased by the VPS35[D620N] mutation (fig. S5, C and D; Table S2). We observed an opposite response for levels of Cathepsin C whose lysosomal levels were markedly increased by the VPS35[D620N] compared to wild type. The levels of Cathepsin B, Cathepsin D, GBA and LAMP1 and TFEB were not altered in VPS35[D620N] LysoTag-IP MEFs. Although the levels of TFE3 transcription factor was moderately increased in the LysoTag-IP of VPS35[D620N] MEFs, it should be noted that TFE3 and TFEB are not strongly enriched in the LysoTag-IP compared to mock and whole cell lysates (fig. S5C and Table S2). These data suggest that transcriptional responses to LRRK2 activity are likely to be cell type specific.

### LRRK2 activity reduces RILPL1 expression in whole cell extracts

MS data of whole cell lysates revealed that levels of RILPL1 were reduced in the VPS35[D620N] background, suggesting that recruitment of RILPL1 to the lysosome accelerates its degradation (Fig. 1, E and F). To explore whether this was linked to LRRK2 kinase activity, we treated VPS35[D620N] knock-in MEFs with MLi-2. Within 8h, expression of RILPL1 returned to levels similar to that of wild type cells (Fig. 3A). Levels of RILPL1 in homozygous VPS35[D620N] mouse brain (Fig. 3B) and lung extracts (Fig. 3C) were also ∼40% lower compared to wild type. Analysis of tissues from VPS35[D620N] mice fed with a diet containing MLi-2 for a 2-week period revealed that inhibiting LRRK2 significantly increased RILPL1 expression in whole cell lysates, ∼1.6-fold in the brain (Fig. 3D) and ∼2.5-fold lung (Fig. 3E).

**Fig 3.**
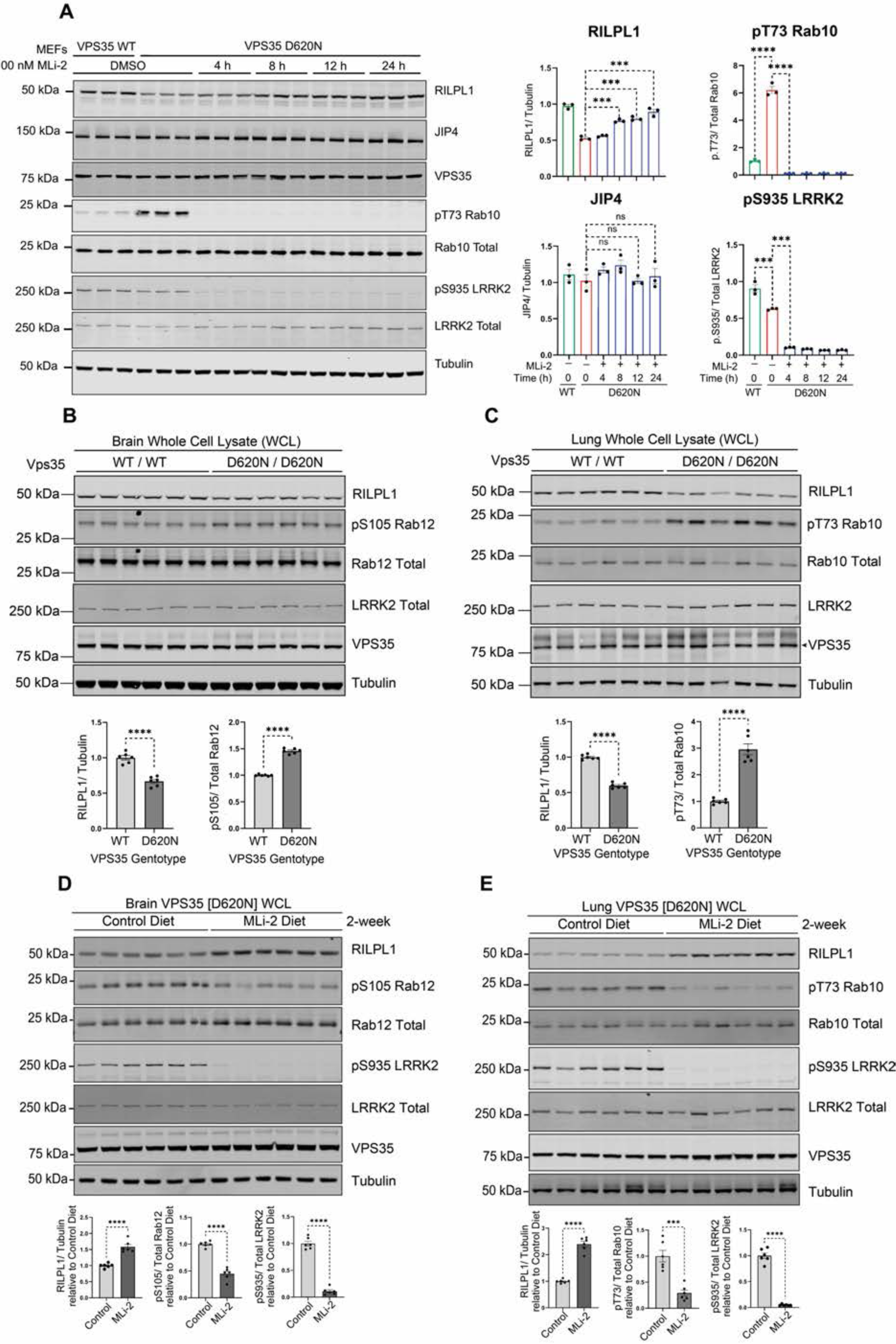
Enhanced LRRK2 activity by VPS35[D620N] mutation reduces expression of RILPL1. (A) Littermate-matched wild type (WT) and VPS35[D620N] homozygous knock-in MEFs were treated ± 100 nM MLi-2 for the indicated times prior to lysis. Lysates were subjected to quantitative immunoblot analysis of the LI-COR Odyssey CLx Western Blot imaging system and indicated antibodies. Technical replicates represent cell extract obtained from a different dish of cells. Quantitation of immunoblotting data (performed using ImageStudioLite software version 5.2.5, RRID:SCR_013715; http://www.licor.com/bio/products/software/image_studio_lite/) is shown as mean ± SEM. Data were analyzed using two-tailed unpaired t-test (** p< 0.01, *** p< 0.001, **** p< 0.001). (B) Brain and (C) lung tissues were harvested from 4-month-old, littermate matched wild type (WT) and homozygous VPS35[D620N] knock-in mice. 20 μg of whole tissue extract were subjected to immunoblot analysis using indicated antibodies as described in (A). (D, E) 4-month-old littermate-matched homozygous Vps35[D620N] knock-in mice were fed on either a control diet or MLi-2 diet for 2-weeks prior to tissue harvesting. Brain (D) and lung (E) tissues extracts were analyzed by immunoblotting and quantification performed as previously described in (A).

### LRRK2 activity drives association of RILPL1 with TMEM55B, a lysosomal integral membrane protein

We hypothesized that in VPS35[D620N] cells, if LRRK2 was recruited to lysosomes and phosphorylated Rab proteins at this location, this could trigger recruitment of RILPL1 to the lysosome, where RILPL1 might interact with other lysosomal protein(s). To explore this further, we undertook a RILPL1 enrichment MS analysis in HEK293 cells overexpressing LRRK2[Y1699C] together with the GTP locked form of Rab8A (Rab8A[Q67L]). As control, we employed a RILPL1 mutant in which the critical Arg293 residue required for binding LRRK2 phosphorylated Rab8A is mutated to Ala; this would block RILPL1 from being recruited to the lysosome by binding to phosphorylated Rab8A (*11*). Control phos-tag immunoblot analysis (*47*), confirmed that in these experiments, Rab8A is phosphorylated by LRRK2 to a significant ∼50% stoichiometry maximizing the opportunity to identify downstream targets (Fig. 4A). Employing an isobaric Tandem Mass Tag (TMT) affinity enrichment MS workflow (Fig. 4B), we compared interactors of wild type RILPL1 with mutant RILPL1[R293A] (Fig. 4C and Table S3). As expected, wild type RILPL1 is associated with Rab8A (fig. S6A) and Rab10 (fig. S6B) to a greater extent than the RILPL1[R293A] mutant (Fig. 4C). In addition, 4 other proteins also interacted more strongly with the wild type RILPL1, namely TMEM55B (Fig. 4C and fig. S6C), a lysosomal integral membrane protein (*48*), Rab34 (Fig. 4C and fig. S6D) (not a known LRRK2 substrate), and two mitochondrial proteins: COX5A (Fig. 4C and fig. S6E) and NDUFA2 (Fig. 4C and fig. S6F).

**Fig 4.**
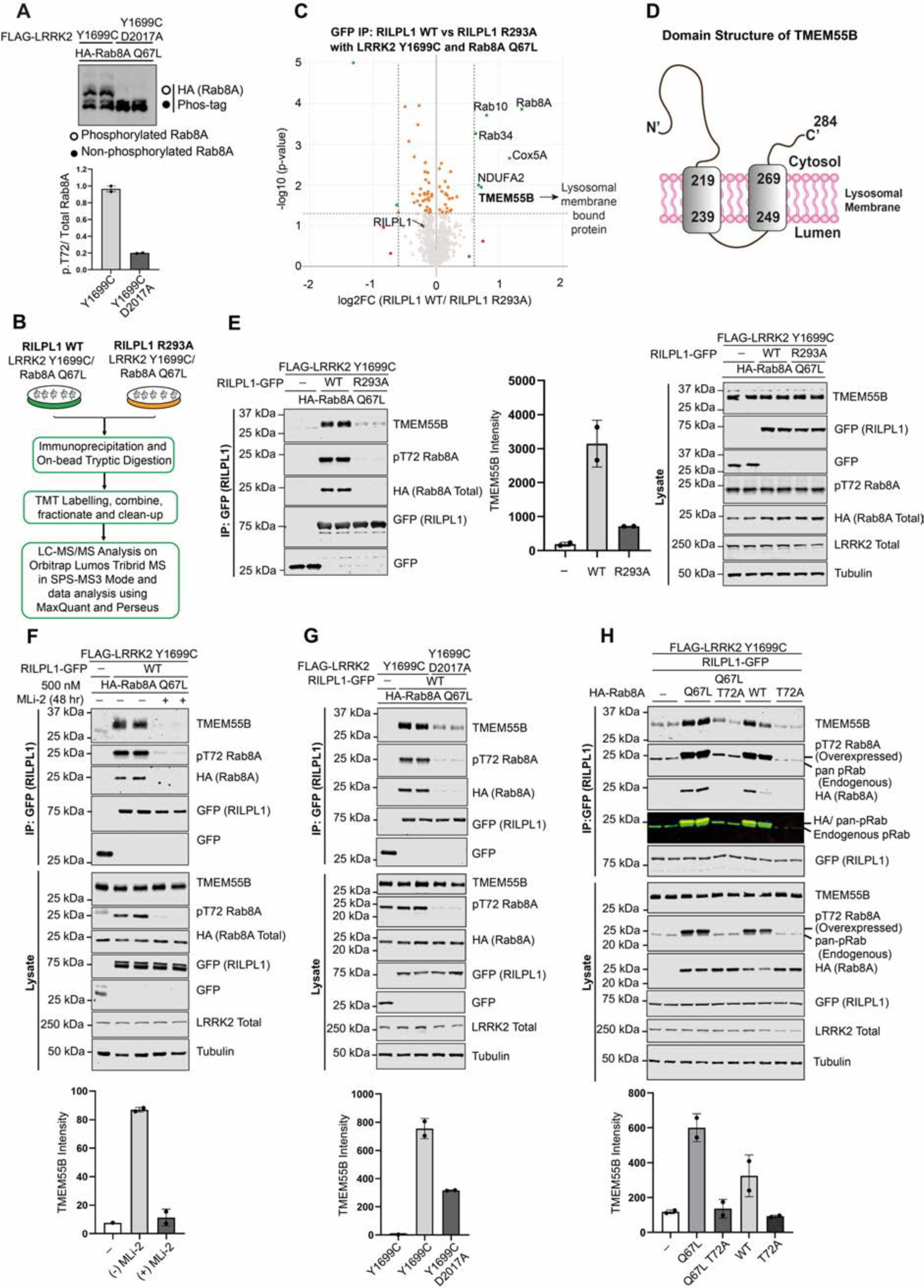
Association of RILPL1 with a lysosomal membrane integral protein, TMEM55B mediated by LRRK2 activity. (A) HEK293 cells were transiently transfected with HA-Rab8A[Q67L] (GTP-bound mutant) either in the presence of LRRK2[Y1699C] (kinase active mutant) or LRRK2[D2017A] (kinase inactive mutant). 24h post transfection 20 µg of whole cell lysate was analyzed on a Phos-tag gel and immunoblot developed using BIO-RAD ChemiDoc Imaging System. Each lane represents cell extracts obtained from a different dish of cells. (B) Depiction of the workflow for the Tandem Mass Tag (TMT) label-based mass spectrometry analysis of GFP-immunoprecipitation (IPs) from HEK293 cells transiently transfected as in (A) with in addition RILPL1-GFP Wild-type (WT) or RILPL1-GFP [R293A] (non pRab8/10 binding mutant) (C) Volcano plot depicting the fold enrichment of proteins between IPs from RILPL1-GFP WT and RILPL1-GFP R293A mutant (*p-*value adjusted by two-tailed *t*-test which is corrected by permutation-based FDR of 5%. (Curtain link: https://curtain.proteo.info/#/4f15d6c8-9192-4bb2-a2ff-da7aa542025c) (Table S3). (D) TMEM55B domain structure. (E to H) Transfections and immunoprecipitation of the indicated proteins were performed as in (B) and analyzed by quantitative immunoblot analysis using the LI-COR Odyssey CLx Western Blot imaging system and indicated antibodies. Quantitation of immunoblotting data (performed using ImageStudioLite software version 5.2.5, RRID:SCR_013715; http://www.licor.com/bio/products/software/image_studio_lite/) is shown as mean ± SEM. Data were analyzed using two-tailed unpaired t-test (** p< 0.01, *** p< 0.001, **** p< 0.001).

We focused on TMEM55B, which is composed of 284 residues and 2 transmembrane domains with N- and C-terminal domains facing the cytosol (Fig. 4D). Immunoblotting experiments validated the MS studies, confirming that endogenous TMEM55B co-immunoprecipitated with wild type, but not mutant RILPL1[R293A] co-expressed with LRRK2[Y1699C] (Fig. 4E). Furthermore, treatment with MLi-2 (Fig. 4F), or introduction of a mutation that ablates LRRK2 kinase activity (D2017A) (Fig. 4G), markedly inhibited association of RILPL1 with TMEM55B. GTP-locked Rab8A[Q67L] associated with TMEM55B to a moderately greater extent than wild type Rab8A (Fig. 4H), consistent with previous data showing that Rab proteins in the GTP-bound conformation interact with higher affinity with RILPL1 (*11*). Taken together these results suggest that when LRRK2 is recruited to VPS35[D620N] lysosomes it phosphorylates Rab proteins which recruit RILPL1 to the lysosome and thereby inducing its interaction with TMEM55B.

### LRRK2 activity promotes co-localization of RILPL1 and TMEM55B at the lysosome

Using confocal microscopy, we investigated the localization of endogenous pRab10, overexpressed Myc-RILPL1 and endogenous TMEM55B in LRRK2[R1441C] MEFs. As shown previously (*13, 14*), we observed significant co-localization of pRab10 and RILPL1 in the perinuclear region that was partially impacted by Nocodazole treatment that disrupts microtubule dynamics (Fig. 5, A and E). Treatment with MLi-2 abolished pRab10 phosphorylation without markedly impacting RILPL1 localization. Some co-localization of Myc-RILPL1 with endogenous TMEM55B was observed in the perinuclear region, and this was also significantly reduced with MLi-2 treatment but not nocodazole (Fig. 5, B and F). To observe the lysosomes in more detail, we utilized expansion microscopy, revealing that a low percentage of lysosomes determined by endogenous TMEM55B expression co-localized with Myc-RILPL1 (Fig. 5, C and G). Treatment with MLi-2 abolished all co-localization of endogenous TMEM55B and Myc-RILPL1 (Fig. 5, C and G). Similar results were obtained in VPS35[D620N] MEFs treated with MLi-2, but not Nocodazole, reducing the interaction of RILPL1 with TMEM55B (Fig. 5, D and H). This data supports the finding that the LRRK2 pathway is driving the association of RILPL1 with TMEM55B in a low proportion of lysosomes.

**Fig 5.**
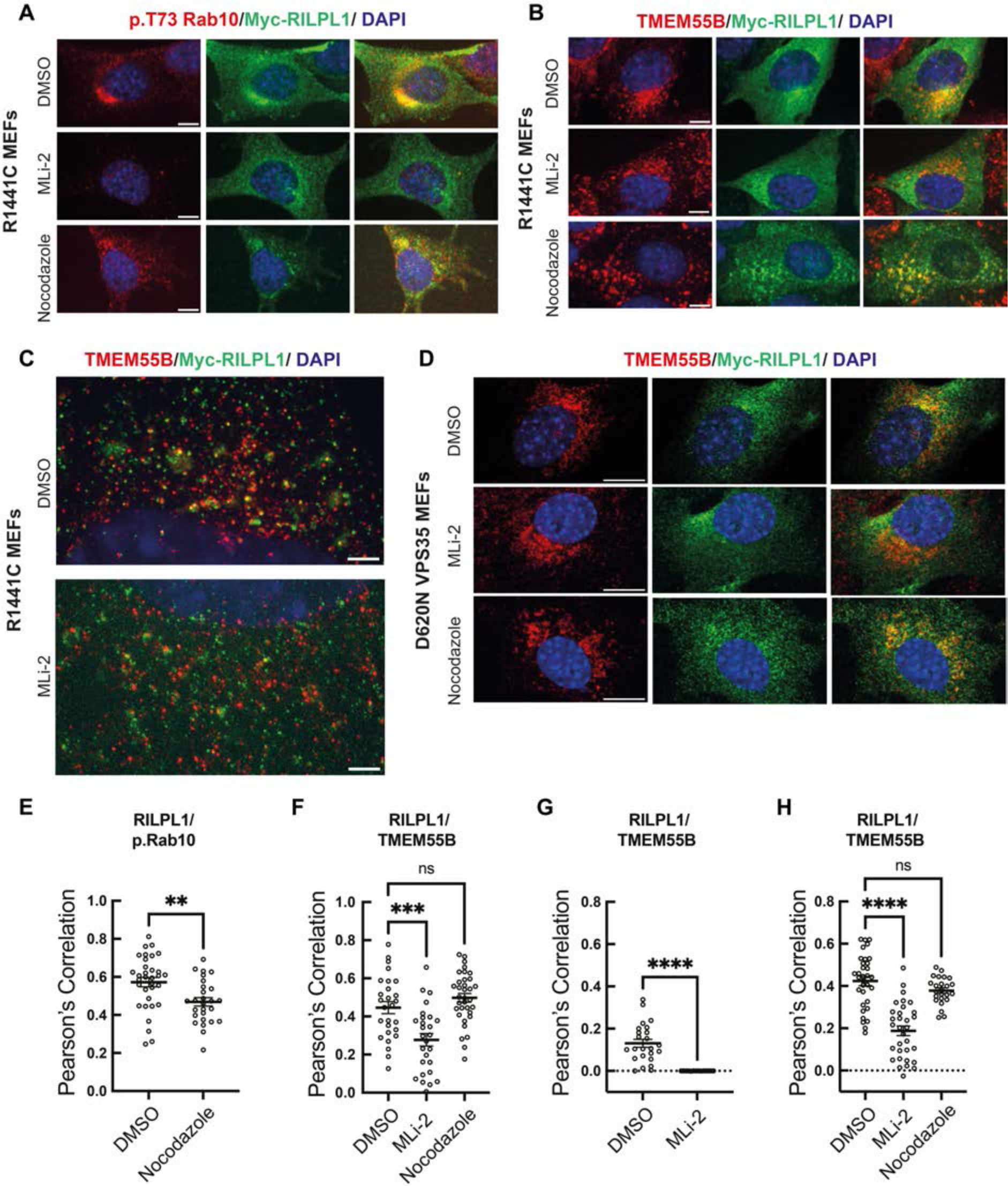
LRRK2-dependent colocalization of RILPL1 and TMEM55B in R1441C and D620N VPS35 MEF cells. Immunofluorescence microscopy of cells expressing transfected Myc-RILPL1 and stained using anti-Myc antibody and antibodies to detect endogenous p.Rab10 or TMEM55B. (A) R1441C MEFs treated with or without 200nM MLi-2 for 2hr or 20µM nocodazole for 2h to depolymerize microtubules. Red, p.Rab10; green, Myc-RILPL1. Merged images are shown at the far right in this and subsequent panels. (B) R1441C MEFs stained to detect endogenous TMEM55B and transfected Myc-RILPL1 as in A. (C) R1441C MEFs visualized using expansion microscopy to detect endogenous TMEM55B (red) and transfected Myc-RILPL1 (green). (D) D620N VPS35 MEFs stained with antibodies to detect endogenous TMEM55B (red) or transfected Myc-RILPL1 (green), and merged image. Scale bars in panels A-D, 10 µm. (E-H) Pearson’s correlation coefficients between RILPL1 and p.Rab10 or TMEM55B as indicated from experiments presented in panels A-D. Significance was determined using Ordinary One-way ANOVA p<0.001 for multiple comparisons or t-test for paired comparisons. (E) **p=0.0032 (F) ***p=0.0003; (G) ****p<0.0001; (H) ****p<0.0001. E, F, and H represent two independent replicates with 25-35 cells total analyzed. G represents one replicate with 15-24 cells analyzed.

### Mapping the RILPL1-TMEM55B interaction interface

Truncation mutagenesis revealed that removal of the last 8 amino acids of RILPL1 (Fig. 6A and fig. S7, A and B), that are evolutionarily conserved (Fig. 6B), abolished binding to TMEM55B. Mutation of several residues within the TMEM-Binding motif (Glu394Lys, Glu398Lys and Ala399Leu) markedly suppressed interaction of RILPL1 with TMEM55B (Fig. 6C). We have termed this region the TMEM55-Binding motif, which is not conserved in RILP and RILPL2. Consistent with this, neither RILP nor RILPL2 co-immunoprecipitate with TMEM55B (fig. S7, C and D).

**Fig 6.**
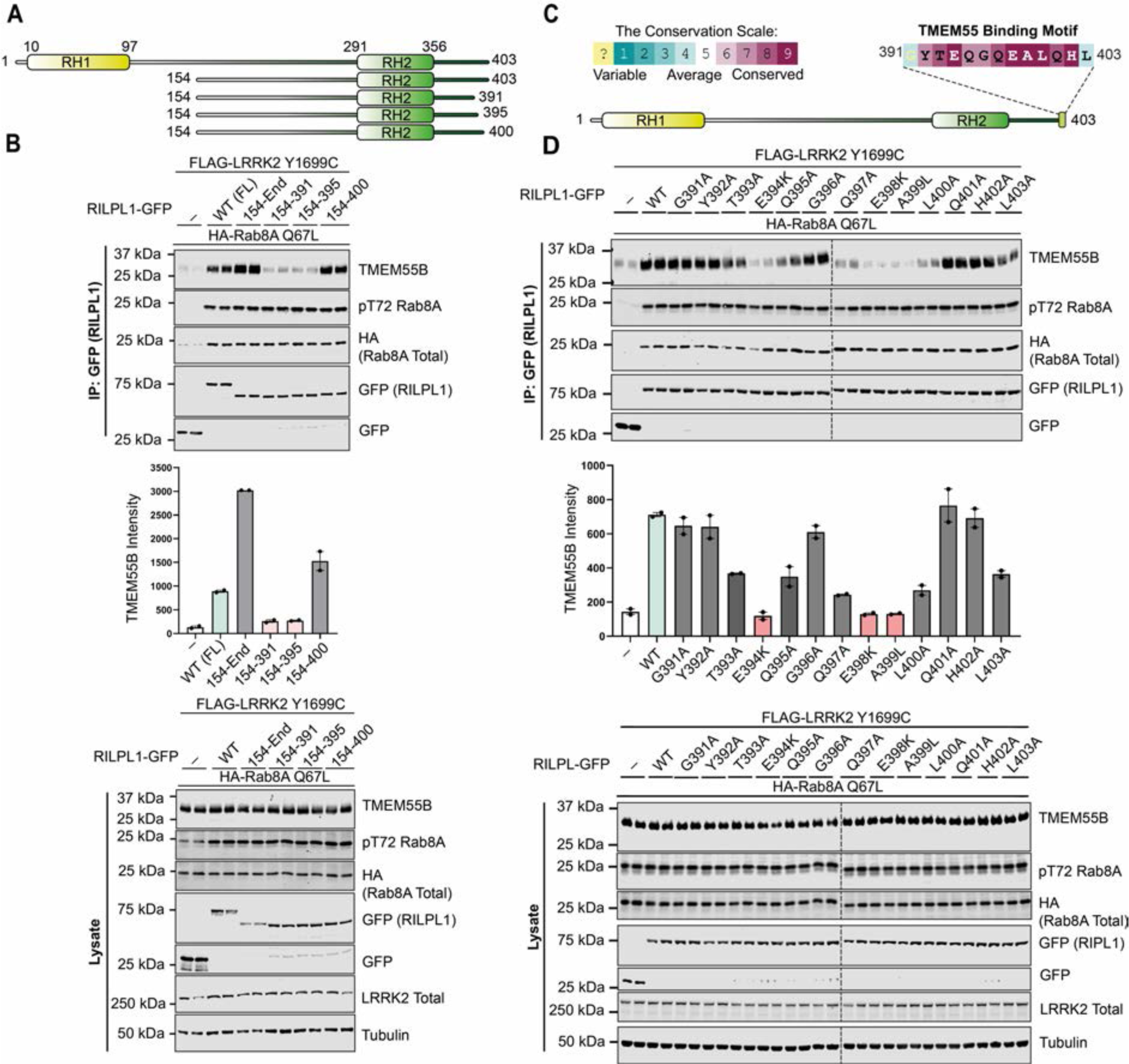
RILPL1 interacts with TMEM55B via a conserved motif at the C-terminal. (A) Domain structure of full length and truncated mutants of RILPL1 used in this study. (B) HEK293 cells were transiently transfected with the indicated proteins and lysed 24h post transfection. GFP-RILPL1 immunoprecipitations (top panel) or cell extracts (lower panel) were subjected to quantitative immunoblot analysis using the LI-COR Odyssey CLx Western Blot imaging system and indicated antibodies. Quantitation of immunoblotting data (performed using ImageStudioLite software version 5.2.5, RRID:SCR_013715; http://www.licor.com/bio/products/software/image_studio_lite/) is shown as mean ± SEM. (C) Analysis of C-terminal residue conservation of RILPL1 using the ConSurf motif software (RRID: SCR_002320) (*64*) and conservation score scale. (D) As in (B).

Mutagenesis analysis revealed that the minimum region of TMEM55B required for interaction with RILPL1 spans an evolutionary conserved region encompassing residues 80-160 (Fig. 7, A and B; fig. S8, A and B) that we have termed the TMEM55-conserved domain. AlphaFold (Jumper et al., 2021) predicts that this region adopts a globular fold possessing a hydrophobic groove along one surface aligned with conserved residues (Fig. 7C). Using AlphaFold2.ipynb ColabFold notebook with AlphaFold2-multimer-v2 and AMBER structure relaxation we modeled how full length RILPL1 would interact with full length TMEM55B. The top models predicted an interaction between the RILPL1 TMEM-Binding-Motif and the hydrophobic groove on the surface of TMEM-Conserved domain, with two additional electrostatic interactions involving conserved residues R151 and K141 (Fig. 7C). Mutational analysis revealed that the R151E but not the K141E mutation ablated binding of TMEM55B to RILPL1 (Fig. 7D). Mutation of the conserved hydrophobic groove residues (V108T, A117S, L137A) also reduced RILPL1 binding (Fig. 7D).

**Fig 7.**
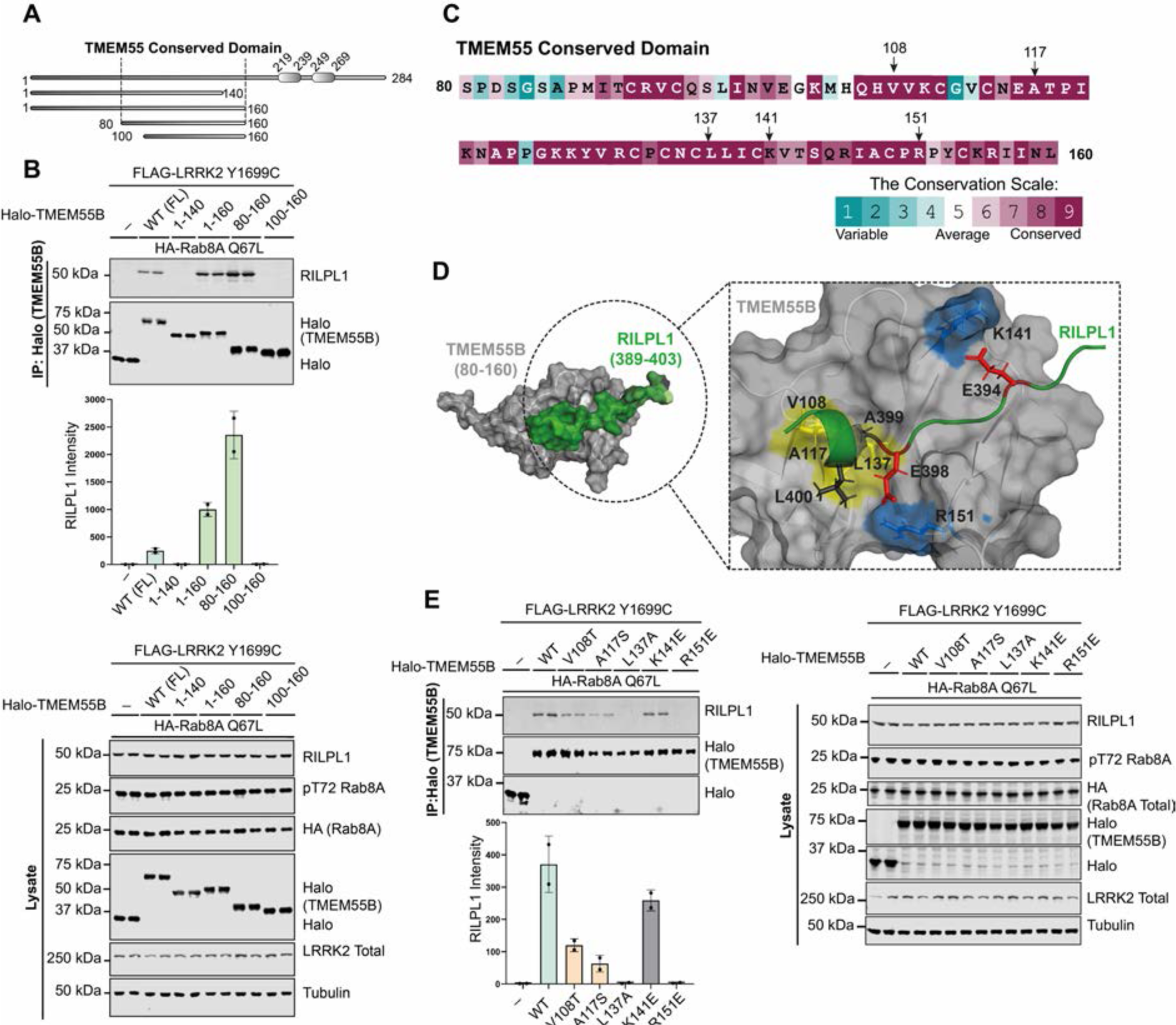
N-terminal conserved domain of TMEM55B facilitates the binding of RILPL1. (A) Domain structure of full length and truncated mutants of TMEM55B used in this study. (B) HEK293 cells were transiently transfected with the indicated proteins and lysed 24h post transfection. Halo-TMEM55B immunoprecipitations (top panel) or cell extracts (lower panel) were subjected to quantitative immunoblot analysis using the LI-COR Odyssey CLx Western Blot imaging system and indicated antibodies. Quantitation of immunoblotting data (performed using ImageStudioLite software version 5.2.5, RRID:SCR_013715; http://www.licor.com/bio/products/software/image_studio_lite/) is shown as mean ± SEM. (C) Analysis of C-terminal residue conservation of TMEM55B using the ConSurf motif software (*64*) and conservation score scale. (D) Key residues which are involved in the interaction between TMEM55B and RILPL1 were analyzed by AlphaFold2 model of TMEM55B [80-160] and RIPL1 [389-403]. (E) As in (B).

Cells express another TMEM55 isoform termed TMEM55A, that is highly related to TMEM55B, with 55% identity in sequence and 86% identity in sequence within the TMEM55 Conserved-domain (fig. S8C). Overexpression studies reveal that TMEM55A, although expressed at ∼2-fold lower levels than TMEM55B, also interacted with RILPL1 (fig. S8D).

### Impact of knockdown of RILPL1 and TMEM55B

We next studied the effect that siRNA knock-down of RILPL1 had on LRRK2 activity and found that this moderately increased LRRK2-mediated phosphorylation of Rab10 and Rab12 in wild-type and VPS35[D620N] MEFs (Fig. 8A). We next observed that siRNA knockdown of TMEM55B, but not TMEM55A, in VPS35[D620N] MEFs increased cellular levels of RILPL1, consistent with the notion that RILPL1 binding to TMEM55B accelerated the degradation of RILPL1 (Fig. 8B). Knockdown of TMEM55A or TMEM55B or both did not impact significantly LRRK2 mediated phosphorylation of Rab10 or Rab12 in VPS35[D620N] MEFs (Fig. 8, B and C). We also generated CRISPR TMEM55B knock-out A549 cells and also observed an increase in RILPL1 levels in whole cell lysate without affecting Rab10 or Rab12 phosphorylation (Fig. 8D).

**Fig 8.**
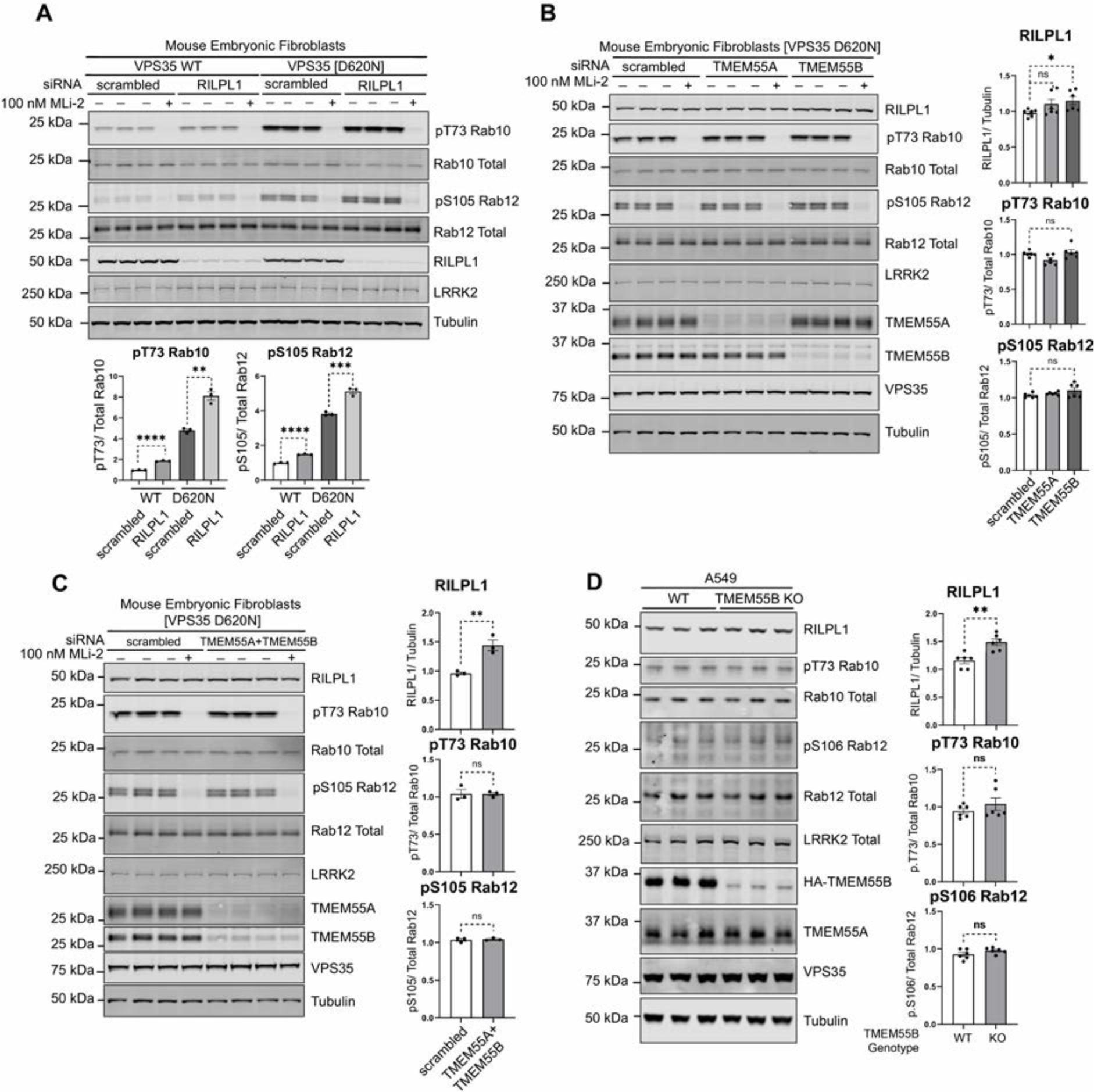
Effect of knockdown or knock-out of RILPL1, TMEM55A and TMEM55B. (A to C) VPS35[D620N] homozygous KI MEFs were transfected with the indicated siRNA for 72 h, and 100 nM MLi-2 (or DMSO, 0.1% v/v) was added 1 h prior to lysis. Cell lysates were analyzed by quantitative immunoblot analysis using the LI-COR Odyssey CLx Western Blot imaging system and the indicated antibodies. Quantitation of immunoblotting data (performed using ImageStudioLite software version 5.2.5, RRID:SCR_013715; http://www.licor.com/bio/products/software/image_studio_lite/) is shown as mean ± SEM. Data were analyzed using two-tailed unpaired t-test (** p< 0.01, *** p< 0.001, **** p< 0.001). (D) TMEM55B CRISPR knock-out (KO) A549 cells and KO cells complemented with virally transduced 3HAxTMEM55B (WT) were lysed and analyzed as in (A).

### The lysosomotropic agent LLOMe also induced recruitment of RILPL1 to the lysosome

Finally, we treated wild type MEFs with the lysosomotropic agent L-leucyl-L-leucine methyl ester (LLOMe) at 1 mM for 1-6 h to determine whether this resulted in the recruitment of RILPL1 to the lysosome. These experiments revealed that LLOMe induced a 1.5 to fold increase in the levels of RILPL1 at the lysosome at later 2-6 h time points (Fig. 9, A and B). In VPS35[D620N] MEFs, LLOMe treatment enhanced lysosomal RILPL1 levels more, namely ∼2.5-fold (Fig. 9, A and B). LLOMe induced a moderate increase in pRab10/ total Rab10 levels in cell lysates (Fig. 9C) and lysosomes (Fig. 9D) that was also ∼2-fold lower than levels caused by the D620N mutation. As observed in Figure 2A, MLi-2 induced a moderate increase in association of LRRK2 to the lysosome, but this was not observed in the absence of MLi-2 with LLOMe treatment of the D620N mutation (Fig. 9E).

**Fig 9.**
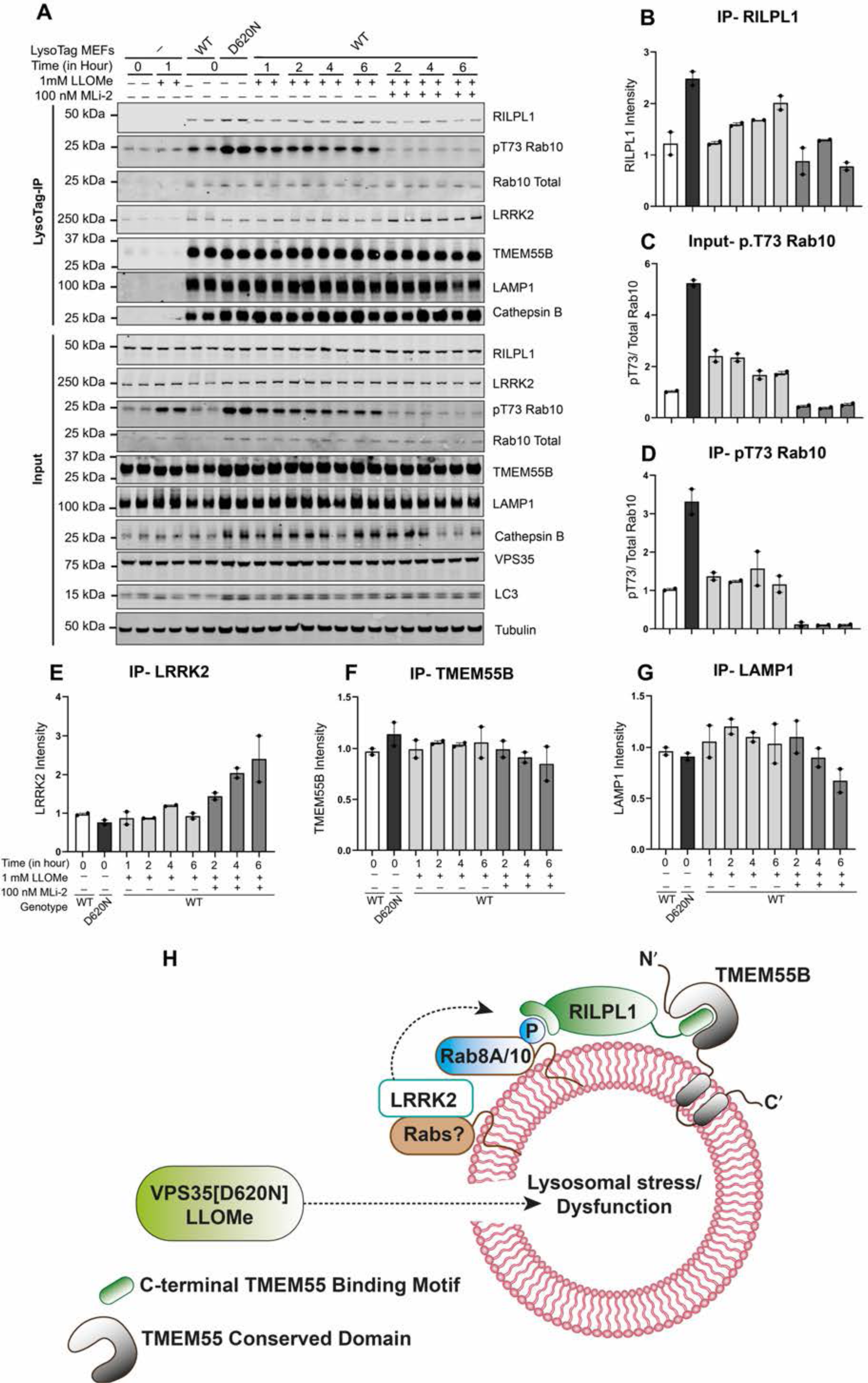
Effect of LLOMe compared to D620N mutation on recruitment of RILPL1 to the lysosome. (A to G) Wild type or VPS35[D620N] homozygous KI MEFs expressing ± LysoTag (TMEM192-3xHA) were treated as indicated ± 1 mM LLOMe and ± 100 nM MLi-2 for the indicated time points. Cells were homogenized and lysosomes immunoprecipitated. 4 μg of whole cell lysate or lysosome extract were analyzed by quantitative immunoblot analysis using the LI-COR Odyssey CLx Western Blot imaging system and indicated antibodies. Quantitation of immunoblotting data (performed using ImageStudioLite software version 5.2.5, RRID:SCR_013715; http://www.licor.com/bio/products/software/image_studio_lite/) is shown as mean ± SEM. (H) Model of how lysosomal dysfunction resulting from VPS35[D620N] mutation recruits and activates LRRK2 to the lysosome, resulting in phosphorylation of Rab proteins which in turn triggers the recruitment of RILPL1 and its binding to TMEM55A/B.

## Discussion

The retromer complex controls retrograde sorting of cargos from the endosome back to the trans Golgi Network, as well as recycling cargo from the endosome to the cell surface. The D260N mutation impacts the core VPS35 backbone subunit of the retromer complex and has been suggested to disrupt the retromer’s retrograde cargo trafficking pathway (*49-51*). Structural analysis of the retromer complex has indicated that the D620N mutation might impact oligomerization of the complex but this has not been definitely established (*52*). Immunoprecipitation MS studies indicate that the D620N mutation moderately impaired interaction with FAM21, a key member of the WASH complex that binds to VPS35 and this mutation thus impairs association of the retromer complex to the endosomes (*49, 50*). How VPS35[D620N] mutations stimulate LRRK2 pathway activity is not known and there is no strong evidence that LRRK2 and the retromer complex interact directly.

Our preferred model is that the D620N mutation, by disrupting the retromer’s retrograde cargo trafficking pathway, ultimately leads to a specific form of lysosomal dysfunction, and this is what triggers LRRK2 activation. Many previous studies have pointed towards lysosomal damage/dysfunction triggering LRRK2 pathway activation. For example, treatment of cells with lysosome-damaging agents such as chloroquine (*53, 54*) or LLOMe (*12*), triggers LRRK2 recruitment to the lysosome inducing its activation and enhanced phosphorylation of Rab8A and Rab10 on the lysosome. Chloroquine was also reported to induce relocalization of Rab8/Rab10 effectors EHBP1 and EHBP1L1 to the lysosome (*53*), whereas LLOMe induced lysosomal recruitment of the pRab10 motor adapter protein JIP4 (*12*) as well as the RAB7 GTPase-activating protein TBC1D15 (*55*). Similarly in macrophages, pathogen infection induces lysosomal damage, and was shown to result in relocalization of LRRK2 to the damaged lysosome where it phosphorylated Rab8A and this recruited the ESCRT-III component CHMP4B that orchestrates the repair of lysosome damage (*30*). A recent study has shown that overexpression of LRRK2 in 293A cells induces perinuclear clustering of lysosomes that display elevated phosphorylation of Rab12 and RILPL1 that are consistent with our findings in this study (*56*).

None of these above studies used VPS35[D620N] as a possible form of more physiological lysosome stress that is relevant to Parkinson’s. We pursued our D620N data for the levels of all of these proteins in wild type and D620N cell lysates and LysoTag-IP (fig. 8S1). The data shows that for CHMP4B (fig. S9A), TBC1D15 (fig. S9B), EHBP1 (fig. S9C) and JIP3 (fig. S9D) levels are moderately decreased in cell lysates of D620N mutation compared to wild type, but there is no clear-cut enrichment of any of these proteins to the lysosome. Treatment of D620N cells with 100 nM MLi-2 for 48h did not alter the levels of these proteins, suggesting that reduced expression in the D620N cells is not related to LRRK2 kinase activity. More work will be required to understand why the D620N mutation moderately reduces the levels of these proteins. For JIP4 (fig. S9E) there were no significant changes or enrichment to the lysosome. We did not detect RILP and RILPL2 in our data sets.

Our data reveals that the VPS35[D620N] mutation in MEFs is indeed having a significant impact on the lysosomal expression of proteins. We observed that the lysosomal abundance of ∼150 proteins was altered over 2-fold in VPS35[D620N] MEFs compared to wild type (Fig 1D). A recent study also reported profound protein abundance changes in LysoTag-IP fractions of H4 neuroglioma VPS35 knock-out cells (*57*). Strikingly, treatment of D620N MEFs with MLi-2, only reduced lysosomal association of a single protein significantly, namely RILPL1. To our knowledge RILPL1 has not been previously observed as a lysosomal associated protein, however previous work was not undertaken in cells expressing elevated LRRK2 activity. Our data also suggests that LLOMe treatment of MEFs VPS35 WT cells induces recruitment of RILPL1 to the lysosome, albeit to a lower extent than observed with D620N mutation (Fig. 9A).

We also find that in homozygous VPS35[D620N] MEFs as well as mouse brain and lung, the levels of RILPL1 in whole cell extracts are reduced compared to wild type and this is largely reversed by treatment with MLi-2. Our model is that once RILPL1 is recruited to the lysosome in VPS35[D620N] cells and tissues, this leads to its accelerated degradation, thereby lowering the steady state level in whole cell lysates. In future work it would be interesting to explore the mechanism by which RILPL1 is degraded at the lysosome and whether reduction of RILPL1 levels in whole cell lysates is a useful biomarker for monitoring lysosomal dysfunction that is of relevance to Parkinson’s disease.

Our data indicates that the recruitment of RILPL1 to TMEM55B is dependent on LRRK2 kinase activity as this is blocked by both MLi-2 and by a mutation that ablates the ability of RILPL1 to bind phospho-Rab proteins. In VPS35[D620N] cells, our data demonstrates that LRRK2 is recruited to the damage/dysfunctional lysosome by a yet unknown mechanism. Using overexpression and immunofluorescence others have demonstrated recruitment of LRRK2 to lysosomes following chloroquine and LLOMe treatment. We and others are unable to detect endogenous LRRK2 by immunofluorescence. However, in our LysoTag-IP methods it is likely that endogenous LRRK2 is dissociated from the lysosome during the cell lysis and/or the washing steps that are involved in the LysoTag-IP steps if it is not strongly anchored to the organelle. One mechanism by which LRRK2 could be recruited to the lysosome involves Rab binding to LRRK2’s N-terminal ARM domain. At the lysosome LRRK2 would phosphorylate Rab proteins including Rab8A and Rab10 either unusually on the lysosome or on adjacent vesicles. This in turn recruits RILPL1 to the lysosome where it then binds to the integral lysosomal transmembrane receptor protein TMEM55B. Our imaging data in Figure 5C suggests that only a low proportion of lysosomes have RILPL1 associated with TMEM55B. It is possible that these represent the actively stressed lysosomes that are being targeted by the LRRK2 pathway. Microscopy studies confirm that MLi-2 treatment ablates binding of RILPL1 and TMEM55B. A model of how binding of RILPL1 and TMEM55 may be regulated is shown on Figure 9H.

The binding mechanism that we have defined for how RILPL1 interacts with TMEM55B involves evolutionary conserved residues on both proteins. Our data suggest that TMEM55A could also be involved in binding to RILPL1 (fig. S8D). The copy number levels of TMEM55A and TMEM55B in HEK293 cells are 30,000-TMEM55A and 141,000-TMEM55B; in MEFs 23,000-TMEM55A and 87,000-TMEM55B and in mouse brain 58,000-TMEM55A and 37,000-TMEM55B (https://copica.proteo.info/#/home). Previous work suggested that TMEM55A and TMEM55B might act as a phosphatidylinositol-4,5-bisphosphate 4-phosphatases based on the presence of a CX5R motif that is located within the TMEM55 Conserved domain (*58*). The AlphaFold model of this domain bears no resemblance to any inositol phosphatase that we are aware of, and it seems likely that this domain functions as a receptor protein to recruit specific proteins such as RILPL1 to the lysosome. Consistent with this another recent study has reported that recombinant TMEM55B lacked detectable inositol phosphatase activity when expressed in vitro (*48*).

Previous work has shown that TMEM55B controls lysosomal movement in cells by binding to the JIP4 motor adaptor protein and thereby linking the lysosome to a dynein-dependent microtubule transport machinery (*48*). It was also reported that TMEM55B levels are transcriptionally upregulated following TFEB and TFE3 activation by starvation or cholesterol-induced lysosomal stress and that this pathway could coordinate lysosome movement in response to stress conditions (*48*). Interestingly, sequence analysis indicates that JIP4 possesses a conserved motif (residues 887 to 898) that is similar in sequence to the C-terminal TMEM binding motif on RILPL1. In future work it would be important to explore whether JIP4 interacts with TMEM55B via this motif, and how these binding interactions might be regulated. Further work is also required to establish how the D620N retromer induced lysosomal stress or LLOMe impacts TMEM55B. We have not observed an increase in TMEM55A (fig. S9F) and TMEM55B (fig. S9G) levels in wild type and D620N MEFs or D620N MEFs treated ± MLi-2. TMEM55B has been reported to be regulated by phosphorylation (*59*) and binding with components of the mTORC1 pathway (*60*). In future work it would be important to understand how TMEM55B is regulated and whether it plays a role in sensing lysosomal dysfunction and what the signaling pathways modulate this process. If TMEM55B and components that it regulates are controlled by lysosomal stress, this could point towards the development of new biomarkers to detect lysosomal stress pathways of relevance to Parkinson’s disease.

## Materials and Methods

### Plasmids

All plasmids used in this study were obtained from the MRC PPU Reagents and Services are available to request via the MRC PPU Reagents and Services website (https://mrcppureagents.dundee.ac.uk). These are listed in Supplementary Materials and Methods.

### Antibodies

All antibodies used in this study are listed in Supplementary Materials and Methods.

### Generation of mouse embryonic fibroblasts (MEFs) and stable expression of LysoTag and GolgiTag

Wild type and homozygous VPS35[D620N] knock-in mouse embryonic fibroblasts (MEFs) were isolated from littermate matched mouse embryos at day E12.5 resulting from crosses between heterozygous mice (Jax strain #: 023409; RRID:IMSR_JAX:023409) using a protocol described in dx.doi.org/10.17504/protocols.io.eq2ly713qlx9/v1. Genotypes were verified via allelic sequencing.

Wild type and homozygous VPS35[D620N] knock-in MEFs stably expressing the LysoTag (*42*) or GolgiTag (*45*) were generated using the protocol described in dx.doi.org/10.17504/protocols.io.6qpvr456bgmk/v1. Briefly, littermate matched primary MEFs were first immortalized by SV40 Large T Antigen viral transduction prior to the stable expression of target proteins (LysoTag, GolgiTag or HA-Empty (vector only encoding a HA tag) by a subsequent viral transduction. To generate desired viruses, HEK293 FRT cells were seeded to give a 60% confluency for transfection the following day. A mixture of 12 μg of DNA (6 μg target DNA, 3.8 μg GAG/POL, 2.2 μg VSVG) and 36 μl of Polyethylenimine (PEI, Polysciences) transfection reagent was diluted to 500 μl in Opti-MEM™ Reduced serum medium (Gibco™) and incubated for 30 minutes prior to adding to cells dropwise. The transfection-medium mixture was discarded after 24h, and 10 ml of fresh medium added. The next day, the medium was harvested and filtered through a 0.45 μm filter before storing at -80°C.

MEF cells were plated in 10 cm dishes to give a 70% confluency the following day for viral transduction by the addition of 5 ml of virus with 5 ml of fresh medium and 10 μg/ml Polybrene (Sigma-Aldrich, TR-1003-G). After 24h incubation, the viral medium was discarded. Cells were given time to recover if required. Cells were selected for viral uptake by addition of a selection agent until non-transduced control plates died. For SV40 Immortalization of MEFs, 200 μM of hygromycin (InvivoGen, ant-hg-5) was used for selection pressure. For Organelle Tag introduction to MEFs, 2 μg/ml puromycin was used for selection pressure.

### Generation of A549 TMEM55B Knock-Out Cells by CRISPR-Cas9 and introduction of 3xHA TMEM55B into A549 TMEM55B Knock-Out Cells

A full transcript map of the TMEM55B locus was constructed by combining data from both NCBI (NC_000014.9) and Ensembl (ENSG00000165782). Knockout (KO) guide RNAs were selected to target the exon 2 to ensure complete disruption of all possible transcripts. Three sets of CRISPR-Cas9 guide RNAs were designed to target exon 2 of TMEM55B: a pair targeting exon 2 (Sense A and Anti-sense A); G1, a single guide RNA (3’-GCCCTTAACTAGCCCGGACAG-5’)’ and G2, a single guide RNA (3’-GACTCGGCAGGTGATCATAG-5’). A549 cells were co-transfected with 1 μg of each plasmid and 2 μg of PEI mixture supplemented with Opti-MEM. 24h post-transfection, cells were kept in DMEM containing 2 µg/ml of Puromycin for 48 h. After the recovery, cell pools were analyzed for the depletion of TMEM55B expression by immunoblotting and afterwards, single cells were sorted using FACS. Following two to three weeks of recovery, promising clones were verified by PCR, shotgun cloning, and sequencing. To rescue the expression of TMEM55B in A549 TMEM55B knock-out cells, we utilized a retrovirus approach to introduce 3xHA-TMEM55B as described in dx.doi.org/10.17504/protocols.io.kqdg3prxpl25/v1.

### Cell culture, transfection, treatments, and lysis

Wild-type and homozygous VPS35[D620N] knock-in MEF cells isolated from littermate-matched mouse embryos were cultured in Dulbecco’s modified Eagle medium (DMEM, Gibco) containing 10% (v/v) foetal bovine serum (FBS, Sigma) 2 mM L-glutamine, 100 U/ml penicillin, and 100 µg/ml streptomycin supplemented with 1X Non-Essential Amino Acid solution and 1 mM sodium pyruvate (Life Technologies, Gibco™). HEK293 cells were purchased from ATCC and maintained in DMEM containing 10% (v/v) FBS, 2 mM L-glutamine, 100 U/ml penicillin, and 100 µg/ml streptomycin. All cells were grown at 37°C temperature with 5% CO2 in a humidified atmosphere and regularly tested for mycoplasma contamination.

Transient transfections were performed in HEK293 cells 24h prior to cell lysis using PEI at around 60-70% confluency. Transfections for co-immunoprecipitation assays were done in 10 cm cell culture dishes using 3 µg of Flag-LRRK2 Y1699C or Flag-LRRK2 Y1699C D2017A as indicated, 1 µg of HA control or HA-Rab8A Q67L and 2 µg of RILPL1-GFP or Halo-TMEM55B cDNA construct per dish diluted in 1 ml of Opti-MEM and 20 µg of PEI mixture and incubated for 30 min before being added to the media. HEK293 cells were treated with 500 nM of MLi-2 inhibitor before the transfections. VPS35[D620N] MEFs were treated with 100 nM of MLi-2 after 24 h of seeding either for 48 h or at different time points (4 h, 8 h, 12 h, and 24 h). Unless otherwise stated, cells were lysed in an ice-cold lysis buffer containing 50 mM Tris-HCl, pH 7.5, 1% (v/v) NP-40 alternative or 1% (v/v) Triton-X100, 10% (v/v) glycerol, 150 mM NaCl, 1 mM sodium orthovanadate, 50 mM sodium fluoride, 10 mM sodium β-glycerophosphate, 5 mM sodium pyrophosphate, 0.1 µg/ml microcystin-LR, and 1 tablet of cOmplete Mini (EDTA-free) protease inhibitor (Merck, 11836170001).

Protein lysates were clarified by centrifugation at 17,000 x *g* for 10 min and were quantified by Bradford assay. Human biological cells (A549/HEK293) were sourced ethically, and their research use was in accord with the terms of the informed consents under an IRB approved protocol. Detailed methods for cell transfection and cell lysis can be found in dx.doi.org/10.17504/protocols.io.bw4bpgsn and dx.doi.org/10.17504/protocols.io.b5jhq4j6.

### Immunofluorescence microscopy

MEF cells were grown in a 6 well plate seeded at 50–60% confluency in 2 ml Dulbecco’s modified Eagle medium (DMEM, Gibco) containing 10% (v/v) foetal bovine serum (FBS, Sigma), 2 mM L-glutamine, 100 U/ml penicillin, and 100 µg/ml streptomycin. After 24 hours, cells were transfected with 2µg plasmid DNA per well in Fugene (3:1 ratio) according to the manufacturer’s instructions. After another 24 hours, cells were trypsinized and plated onto 12 mm glass coverslips (Fisher Scientific, US) in a 6 well plate (Fisher Scientific, US) at 60% confluency. After another 24 hours, cells were treated with MLi-2 or nocodazole before processing for microscopy according to this method dx.doi.org/10.17504/protocols.io.j8nlkw1n5l5r/v1. For expansion microscopy, details can be found at dx.doi.org/10.17504/protocols.io.ewov1o8m7lr2/v1.

All images were obtained using a spinning disk confocal microscope (Yokogawa) with an electron multiplying charge coupled device (EMCCD) camera (Andor, UK) and a 63X oil immersion objective or a Zeiss LSM 900 microscope acquired using Zen 3.4 and a 63X objective. Images were converted to maximum intensity projections using Fiji (https://fiji.sc/). For quantitation using CellProfiler (RRID:SCR_007358; http://cellprofiler.org), image maximum intensity projections were processed in batch for using this macro: dx.doi.org/10.17504/protocols.io.3byl4bpo8vo5/v1. Pearson’s correlation coefficients were obtained as described in dx.doi.org/10.17504/protocols.io.rm7vzbqp5vx1/v1.

### siRNA-mediated knockdown of target proteins in MEFs

For siRNA knockdown of proteins of interest, ON-TARGETplus Mouse Tmem55A siRNA-SMARTpool (cat# L-059670-01-0005), ON-TARGETplus Mouse Tmem55B siRNA-SMARTpool (cat# L-047594-01-0005), ON-TARGETplus Mouse Rilpl1 siRNA-SMARTpool (cat# L-063225-01-0005) and ON-TARGETplus non-targeting pool (#D-001810-10-05) were purchased from Horizon Discovery Ltd. MEF cells were seeded in a six-well format at 200,000 cells/well for transfection the following day (at 60-70% confluency). Cells were transfected using Lipofectamine™ RNAiMAX Transfection Reagent (Invitrogen™, cat# 13778075) according to the manufacturer’s protocol (https://assets.thermofisher.com/TFS-Assets/LSG/manuals/Lipofectamine_RNAiMAX_Reag_protocol.pdf). Briefly, 50 pmol siRNA was diluted in 150 μl Opti-MEM and combined with 10 μl Lipofectamine RNAiMAX in 150 μl Opti-MEM per well. The two mixtures were incubated together at room temperature for 5 min and 250 µl was added dropwise to cells, which were harvested 72 h after transfection.

### Mouse models and MLi-2 diet study

All animal studies were ethically reviewed and carried out in accordance with the Animals (Scientific Procedures) Act 1986 and regulations set by the University of Dundee and the U.K. Home Office, and the GSK Policy on the Care, Welfare and Treatment of Animals. Animal studies and breeding were approved by the University of Dundee ethical committee and performed under a U.K. Home Office project licence. Mice were housed at an ambient temperature (20–24°C) and humidity (45– 55%) and were maintained on a 12 h light/12 h dark cycle, with free access to food and water. VPS35[D620N] knock-in mice (Jax strain #: 023409; RRID:IMSR_JAX:023409) crossed with LysoTag knock-in mice (Jax strain #: 035401; RRID:IMSR_JAX:035401) were used for this study. Mouse genotyping was performed by PCR using genomic DNA isolated from tail clips or ear biopsies. For the experiment shown in Figure 3D and 3E, littermate or age-matched male and female VPS35[D620N] homozygous knock-in mice at 4 months of age were used. Mice were allowed to acclimatize to the control rodent diet (Research Diets D01060501; Research Diets, New Brunswick, NJ) for 14 days before being placed on study. On day 1 of the study, one group (6 mice) received modified rodent diet (Research Diets D01060501) containing MLi-2 and formulated by Research Diets to provide a concentration of 60 mg/kg per day on the basis of an average food intake of 5 g/day for 14 days; the other group (12 mice) received untreated diet (Research Diets D01060501) for 14 days and served as the control group. The dose of MLi-2 and the length of the in-diet treatment used for this study were based on (*46*). Bodyweight and food intake were assessed twice weekly. On the last day of the study, all mice were euthanized by cervical dislocation and the brain and lung tissues were transferred to ice-cold PBS and processed immediately as described below.

### Mouse tissue lysis for organelle immunoprecipitation

The brain and lung tissue were transferred to a cold room and subjected to homogenization using a 2 ml Dounce homogeniser (VWR, Tissue grinders, Potter-Elvehjem type, 432-0206) by 25 strokes in 1 ml of KPBS (136 mM KCl, 10 mM KH2PO4, pH 7.2 using KOH) supplemented with 1X cOmplete Mini (EDTA-free) protease inhibitor (Merck, 11836170001) and 1X PhosSTOP™ phosphatase inhibitor (Merck, 4906837001). The homogenate was collected and precleared by centrifugation at 1,000 x *g* for 2 minutes at 4°C. The precleared homogenate was collected and lysed in a 1:1 dilution with a ice-cold lysis buffer as stated above except the detergent used was 1% Triton X-100 (v/v) instead of 1% NP-40 (v/v). The lysate was kept on ice for 10 minutes before clarification by centrifugation at 17,000 x *g* for 10 minutes at 4°C followed by protein concentration estimation and immunoblot analysis as stated above. A detailed method for isolation of organelles from mouse tissues is described in dx.doi.org/10.17504/protocols.io.x54v9d61zg3e/v1. Isolated organelles were processed as described in dx.doi.org/10.17504/protocols.io.ewov1o627lr2/v1.

### Organelle isolation from MEFs

Lysosomes were isolated from wild-type and homozygous VPS35[D620N] knock-in MEF cells stably expressing the LysoTag (TMEM192-3xHA) via HA immunoprecipitation. MEFs cultured to confluency in 15 cm cell culture dishes were washed briefly with PBS prior to cell scraping into 1 ml KPBS buffer. Cells were pelleted at 1500 x g for 2 mins at 4°C. The KPBS supernatant was then aspirated and 1 ml fresh KPBS was used to resuspend the cell pellet. For downstream whole cell lysate analysis, 50 µl of this cell suspension was retained and lysed in 1% Triton X-100 (v/v) lysis buffer as described above. The remaining cell suspension (950 µl) was subjected to ball-bearing homogenization with an Isobiotec cell homogenizer with 10 micron clearance, involving 10 passes back and forth of the sample through the ball-bearing device. The homogenized cell sample was recovered and centrifuged (1500 x g for 2 mins at 4°C). The supernatant was then applied to 100 µl anti-HA Pierce magnetic beads (Thermo Fisher Scientific) in a 1.5 ml Eppendorf, mixing gently via pipetting up and down 5 times. The sample tube was then placed on an IBI Scientific belly dance orbital shaker, set to full speed, at 4°C for 5 min. Following the 5 min IP incubation, IP tubes were placed on a magnetic tube holder for 30 seconds, before the supernatant was removed. The magnetic beads, with bound lysosomes, were then washed 3 x 1 ml in KPBS buffer (transferring the beads to a fresh tube at the 3rd wash), using the magnetic sample holder to draw beads from solution during each wash. After the final KPBS wash, the bead sample was either lysed directly in 1% Triton X-100 lysis buffer for immunoblotting analysis or stored dry at -80°C prior to sample preparation for proteomic analysis. A detailed method for organelle isolation and analysis is described in dx.doi.org/10.17504/protocols.io.ewov1o627lr2/v1.

### Co-immunoprecipitation assays

GFP or Halo immunoprecipitation were performed according to the manufacturer’s protocol (For GFP IP: GFP-Trap Agarose - ChromoTek GmbH, for Halo IP: https://www.promega.co.uk/-/media/files/resources/protocols/technical-manuals/0/halolink-resin-protocol.pdf) as described in dx.doi.org/10.17504/protocols.io.eq2ly7kxqlx9/v1. Briefly, lysates were incubated with either GFP-Trap agarose beads (Chromotek) or HaloLink Resin (Promega) for 1-2 h (20 µl of packed resin/ 1 mg of lysate). Immunoprecipitates were washed three times with wash-buffer (50 mM Tris-HCl pH 7.5, 150 mM NaCl) and eluted by adding 2x NuPAGE LDS sample buffer. The mixture was then incubated at 95°C for 10 min and the eluent was collected by centrifugation through a 0.22 µm Spin-X column (CLS8161, Sigma). Eluted samples were supplemented with 1% (by volume) β-mercaptoethanol and denatured at 70°C for 10 min before being subjected to immunoblot analysis.

### Quantitative Immunoblotting Analysis

Quantitative immunoblotting analysis was performed according to the protocol described in dx.doi.org/10.17504/protocols.io.bsgrnbv6. Briefly, 10-20 µg of lysate or 25% of the immunoprecipitated samples were loaded onto NuPAGE 4–12% Bis–Tris Midi Gels (Thermo Fisher Scientific, Cat no. WG1402BOX or Cat no. WG1403BOX) or self-cast 10% Bis–Tris gels (0.375 M Bis-tris pH 6.8, 10% w/v acrylamide, 1% v/v TEMED and 0.05% w/v APS) and electrophoresed at 130 V for 2 h with NuPAGE MOPS SDS running buffer (Thermo Fisher Scientific, Cat no. NP0001-02). At the end of electrophoresis, proteins were electrophoretically transferred onto a nitrocellulose membrane (GE Healthcare, Amersham Protran Supported 0.45 µm NC) at 90 V for 90 min on ice in transfer buffer (48 mM Tris and 39 mM glycine supplemented with 20% (v/v) methanol). The membranes were blocked with 5% (w/v) skim milk powder dissolved in TBS-T (50 mM Tris base, 150 mM sodium chloride (NaCl), 0.1% (v/v) Tween 20) at room temperature for 1 h. Membranes were washed three times with TBS-T and were incubated in primary antibody overnight at 4°C. Prior to secondary antibody incubation, membranes were washed three times for 15 min each with TBS-T. The membranes were incubated with secondary antibody for 1 h at room temperature. Thereafter, membranes were washed with TBS-T three times with a 15 min incubation for each wash, and protein bands were acquired via near-infrared fluorescent detection using the Odyssey CLx imaging system and intensities of bands quantified using Image Studio Lite (Version 5.2.5, RRID:SCR_013715).

For Phos-tag analysis, samples were supplemented with 10 mM MnCl2 before loading them onto the gel. Phos-tag gel consisted of stacking gel [4% w/v acrylamide, 0.125 M Tris-HCl, pH 6.8; 0.2% v/v Tetramethylethylenediamine (TEMED) and 0.08% w/v ammonium persulphate (APS)] and resolving gel [10% w/v acrylamide, 375 mM Tris-HCl, pH 8.8; 75 µM PhosTag reagent (MRC PPU Reagents and Services), 150 µM MnCl2, 1% v/v TEMED and 0.05% w/v APS]. Samples were loaded onto the gel after centrifugation at 17,000 x *g* for 1 min and electrophoresed at 90 V with running buffer [25 mM Tris-HCl, 192 mM glycine, and 0.1% (w/v) SDS]. For immunoblot analysis, gels were washed 3×10 min with 48 mM Tris-HCl, 39 mM glycine, 10 mM EDTA, and 0.05% (w/v) SDS followed by one wash with 48 mM Tris-HCl, 39 mM glycine, and 0.05% (w/v) SDS for 10 min. Proteins were then transferred onto the nitrocellulose membranes at 100 V for 180 min on ice using transfer buffer as mentioned before. Membranes were blocked with 5% (w/v) skimmed milk dissolved in TBS-T at room temperature. Next the membranes were incubated with the primary antibodies overnight at 4 °C. After washing the membrane with TBS-T (3×10 min), membranes were incubated with horseradish peroxidase (HRP) conjugated secondary antibody diluted in 5% skimmed milk in TBS-T at room temperature for 1 h. After washing the membranes in TBS-T (5×10 min), protein bands were developed using the ChemiDoc system (Bio-Rad) after adding the ECL reagent (SuperSignal West Dura Extended Duration, Thermo Fisher Scientific) to the membranes.

### Sample preparation, Labeling, Fractionation, LC-MS/MS and data analysis for Tandem Mass Tag (TMT) experiments

The washed GFP immunoprecipitation beads were dissolved in a 100 µl buffer containing 2 M Urea, 50 mM Tris-HCl pH 7.5, 1 mM Dithiothreitol (DTT) incubated on a Thermomixer at 32 °C for 30 minutes and then supplemented with final 20 mM Iodoacetamide(IAA) for another 30 minutes in dark. 250 ng of Sequencing grade trypsin was added to the samples and incubated on a Thermomixer at 1200 rpm agitation for 2Hrs and the supernatant was transferred to new 15.ml Eppendorf tubes and the tryptic digestion was continued for 12h. The reaction was quenched by adding final 1% (v/v)Trifluoroacetic acid and peptides were purified using in-house prepared strong cation exchange stage-tips. Eluted peptides were vacuum dried and tandem mass tags labeling was performed (11-plex TMT, Thermo Scientific) by following manufacturer instructions. Post labeling verification, samples were pooled to equal volumes and vacuum dried. To improve the coverage pooled TMT labeled mix was subjected to mini-basic reversed-phase liquid chromatography fractionation as described in (*61*) and generated a total of four fractions which are vacuum dried and stored at -80°C until LC-MS/MS analysis.

Each fraction was analyzed on a Thermo Orbitrap Lumos Tribrid mass spectrometer in a data dependent (DDA) MS3 mode. The peptides were loaded on a 2 cm pre-column and resolved on a 50 cm analytical column at 300 nl/min flow rate. The full scan was acquired at 120,000 m/z resolution in the mass range of 375-1500 m/z and measured using Orbitrap mass analyzer. The top 10 data dependent MS2 scans were isolated by setting quadrupole mass filter at 0.7 Da and fragmented using 35% collisional induced dissociation. The fragment ions were measured using Ion-trap in a rapid scan mode. Synchronous precursor selection (MS3) for top 10 fragment ions in the mass range of 400 - 1200 m/z were isolated and fragmented using 65% Higher energy collisional dissociation (HCD) and measured at 50,000 m/z 200 resolution using Orbitrap mass analyzer. The Automatic gain control (AGC) targets were set at 2E5, 2E4 and 5E4 for MS1, MS2 and MS3 scans respectively with an ion injection times set at 50 ms for MS1 and MS2 and 120 ms for MS3 scans. The raw MS data was processed using MaxQuant software suite (*62*) (RRID:SCR_014485, version 1.6.6.0; https://www.maxquant.org/). The data type was set as a reporter ion MS3. The data was searched against Human Uniprot (version 2017) by selecting the default contaminants. Carbamdiomethlation of Cys was used as a static modification and Oxidation (M);Acetyl (Protein N-term);Deamidation (NQ); Phosphorylation (STY) were set as variable modifications. 1% FDR was applied at PSM and protein levels. The protein group.txt files were then further processed using Perseus software suite (*63*) (RRID:SCR_015753, version 1.6.0.15; https://www.maxquant.org/perseus/) for statistical analysis. A detailed protocol can be found at dx.doi.org/10.17504/protocols.io.eq2ly7kxqlx9/v1.

### Sample preparation, LC-MS/MS, and data analysis for Data-Independent Acquisition (DIA) experiments

To reduce the lysates, 5 mM of tris(2-carboxyethyl)phosphine (TCEP) was used. The samples were placed on a thermomixer (Eppendorf, UK) at 60°C with 1,100 rpm for 30 min. After cooling to room temperature, 20 mM iodoacetamide (IAA) was added for alkylation. During alkylation, the samples were shielded from light and placed on a thermomixer at 25°C with 1,100 rpm for 30 min. Each sample was then mixed with 5% (v/v) sodium dodecyl sulphate (SDS) and 1.2% (v/v) phosphoric acid, and further diluted with 6X wash buffer (90% MeOH, 10% TEABC at pH 7.2). The samples were thoroughly vortexed and loaded onto S-Trap^TM^ (ProtiFi, USA) columns by centrifugation at 1000 x *g* for 1 min, and the flow-through collected from the columns was discarded. After sample loading, the S-Trap^TM^ columns were washed three times with 150 µL wash buffer. On-column digestion was performed by incubating 60 µL (1.5 µg) Trypsin/Lys-C Mix (Mass Spec Grade; Promega, UK) in 50 mM TEABC solution at pH 8 on the samples on a thermomixer at 47°C for 1 hour before reducing the incubation temperature to 22°C for overnight digestion. The samples were then eluted into 1.5 mL low binding tubes (Eppendorf, UK) by centrifugation with 60 µL of 50 mM TEABC solution at pH 8, followed by 60 µL of 0.15% (v/v) formic acid (FA) aqueous solution, and then 60 µL of elution buffer (80% ACN with 0.15% FA in aqueous solution) twice. The eluted samples were immediately snap-frozen on dry ice and dried at 35°C using a SpeedVac Vacuum Concentrator (Thermo Fisher Scientific, UK). The dried samples were resuspended in 60 µL solution containing 3% (v/v) ACN and 0.1% (v/v) FA, and further incubated on a thermomixer at 22°C with 1200 rpm for 30 min followed by 30 min sonication in a water bath. The sample concentration was then estimated using a NanoDrop instrument (Thermo Fisher Scientific, UK) by measuring the solution absorbance at 224 nm wavelength.

Liquid chromatography tandem mass spectrometry (LC-MS/MS) was performed using an UltiMate 3000 RSLC nano-HPLC system (Thermo Fisher Scientific, UK) coupled to an Orbitrap Exploris^TM^ 480 mass spectrometer (Thermo Fisher Scientific, UK). 4 µg of each sample were loaded onto the nano-HPLC system individually. Peptides were trapped by a precolumn (Acclaim PepMap^TM^ 100, C18, 100 µm x 2 cm, 5 µm, 100 Å) using an aqueous solution containing 0.1% (v/v) TFA. The peptides were then separated by an analytical column (PepMap^TM^ RSLC C18, 75 µm x 50 cm, 2 µm, 100 Å) at 45°C using a linear gradient of 8 to 25% solvent B (an 80% ACN and 0.1% FA solution) for 98 minutes, 25 to 37% solvent B for 15 minutes, 37 to 95% solvent B for 2 minutes, 95% solvent B for 8.5 minutes, 95% to 3% solvent B for 0.5 minutes, and 3% solvent B for 9.5 minutes. The flow rate was set at 250 nL/min for all experiments. Data were acquired in data-independent acquisition (DIA) mode containing 45 isolated m/z windows ranging from 350 to 1500. Collision-induced dissociation (CID) with nitrogen gas was used for peptide fragmentation.

The DIA MS experiment’s raw data were analyzed using the DIA-NN software (RRID: SCR_022865, version 1.8) (*43*) employing a library-free search mode. Trypsin/P was selected as the digestive enzyme, and up to 2 missed cleavages were allowed. Carbamidomethylation at Cysteine residue was set as a fixed modification, while oxidation at methionine residue was included as a variable modification. The software automatically detected and adjusted the mass error (ppm). A protein identification cut-off of 1% FDR was used, and a protein quantification required a minimum of 2 peptides in 5 out of 6 samples. The search results were then imported into Perseus software (*63*) (RRID:SCR_015753, version 1.6.0.15; https://www.maxquant.org/perseus/) for statistical analysis. For the LysoTag-IP samples, IP samples were first compared against the relevant mock IP samples to classify proteins significantly enriched at the lysosome, using a fold-change > 1.5 and p-value < 0.05 (supplementary Figure 1). The lysosomal enriched proteins were then compared against genotypes or treatments to investigate protein level changes at the lysosome organelle. For the whole cell lysate samples, proteins were directly compared against genotypes or treatments to determine the proteome changes in the cells. Significant up-/down-regulated proteins (fold-change > |1.5| and p-value < 0.05) obtained from LysoTag IP and whole cell lysate samples of MEFs VPS35 wild-type versus D620N were then submitted to metascape (RRID:SCR_016620, version 5.3) (*44*) for enrichment analysis. Enrichment of GO biological processes pathway, GO molecular functions, and GO cellular components with p-value < 0.01 were reported in supplementary Figure 3. The text files generated from Perseus software were then imported into an in-house software, Curtain 2.0, for data visualization. A detailed protocol can be found at dx.doi.org/10.17504/protocols.io.ewov1o627lr2/v1.

## Statistical Analysis

All statistical analyses were performed in GraphPad Prism (RRID:SCR_002798, version 9.3.1; http://graphpad.com/). Two-tailed unpaired *t*-test was performed for statistical comparison of two groups.

## Acknowledgements

We thank the support of Monther Abu-Remaileh and his laboratory for help in setting up the LysoTag-IP method and how to best analyze results, as well as Amir Khan for helpful discussion on the modeling of RILPL1-TMEM55B structures. We thank the excellent technical support of the MRC Protein Phosphorylation and Ubiquitylation Unit (PPU) Genotyping team (coordinated by Gail Gilmour), the MRC-PPU DNA sequencing service (coordinated by Gary Hunter), the MRC-PPU tissue culture team (coordinated by Dr Edwin Allen), the MRC-PPU mass spectrometry facility team (coordinated by Dr Renata Soares) and the MRC-PPU Reagents and Services team (coordinated by Dr James Hastie).

## Author contributions

PP Conceptualization Methodology Investigation Writing—original draft Writing— review & editing

MT Conceptualization Methodology Investigation Writing—original draft Writing— review & editing

PYL Conceptualization Methodology Investigation Writing—original draft Writing— review & editing

FT Methodology Investigation Writing—review & editing

CAH Methodology Investigation

PL Methodology Investigation

RSN Methodology Investigation

TKP Methodology

EAD Methodology Investigation

MW Methodology

TM Methodology

SRP Conceptualization Supervision Writing—review & editing

DRA Conceptualization Supervision Writing—original draft Writing—review & editing

## Data and materials availability

All the primary data that is presented in this study has been deposited in publicly accessible repositories. Immunoblotting data and confocal microscopy data (raw data files, their quantitation and statistical analysis) have been deposited in Zenodo (10.5281/zenodo.7886705 and 10.5281/zenodo.7765022). Proteomic data have been deposited in the ProteomeXchange PRIDE repository (Identifier:PXD042502). All plasmids and antibodies generated at the MRC Protein Phosphorylation and Ubiquitylation Unit at the University of Dundee can be requested through our website https://mrcppureagents.dundee.ac.uk/. For the purpose of open access, the authors have applied a CC BY public copyright license to all Author Accepted Manuscripts arising from this submission.

## Supplementary section

### Supplementary Materials and Methods Plasmids

#### Plasmids

All plasmids used in this study were obtained from the MRC PPU Reagents and Services are available to request via the MRC PPU Reagents and Services website (https://mrcppureagents.dundee.ac.uk).

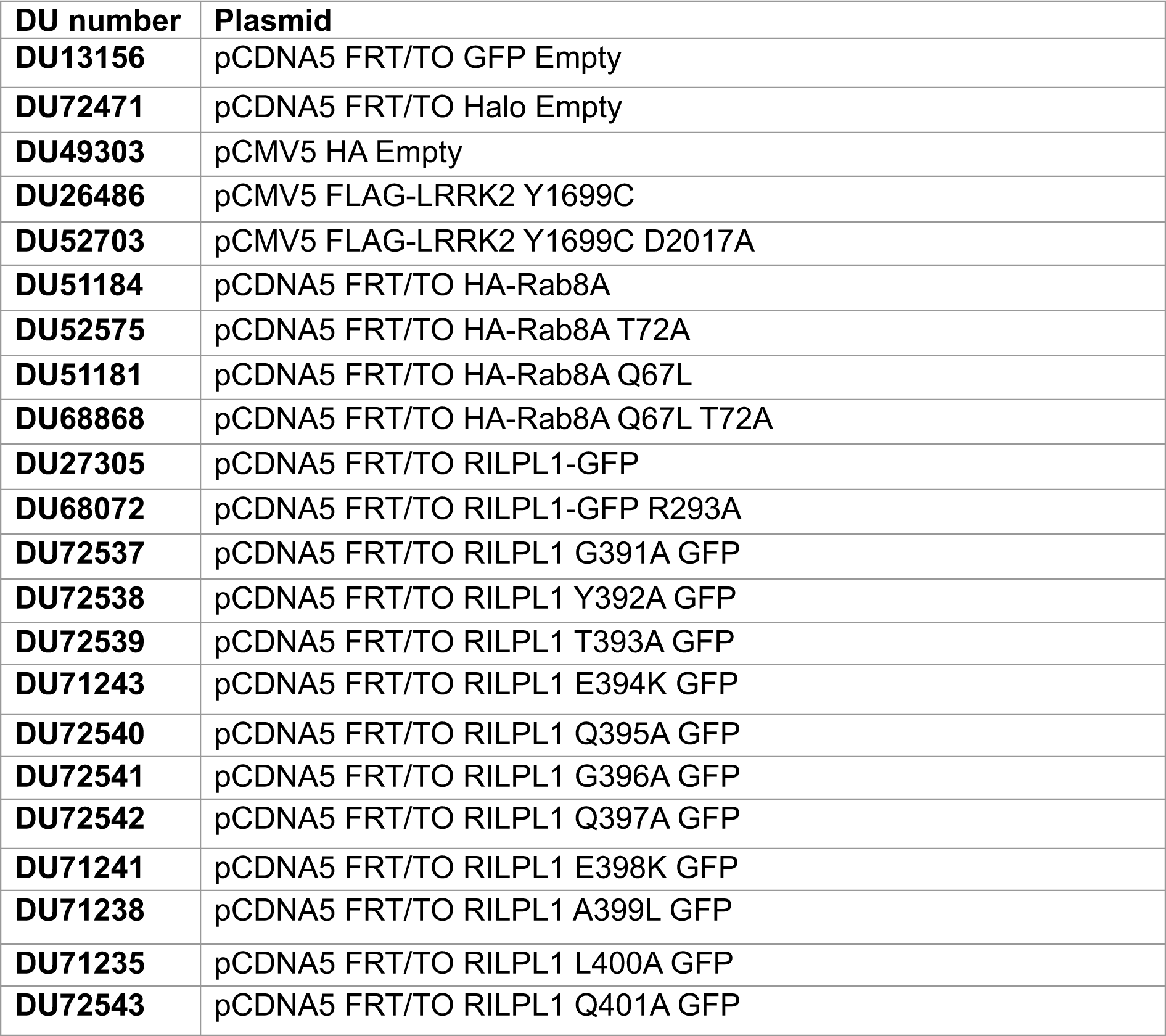
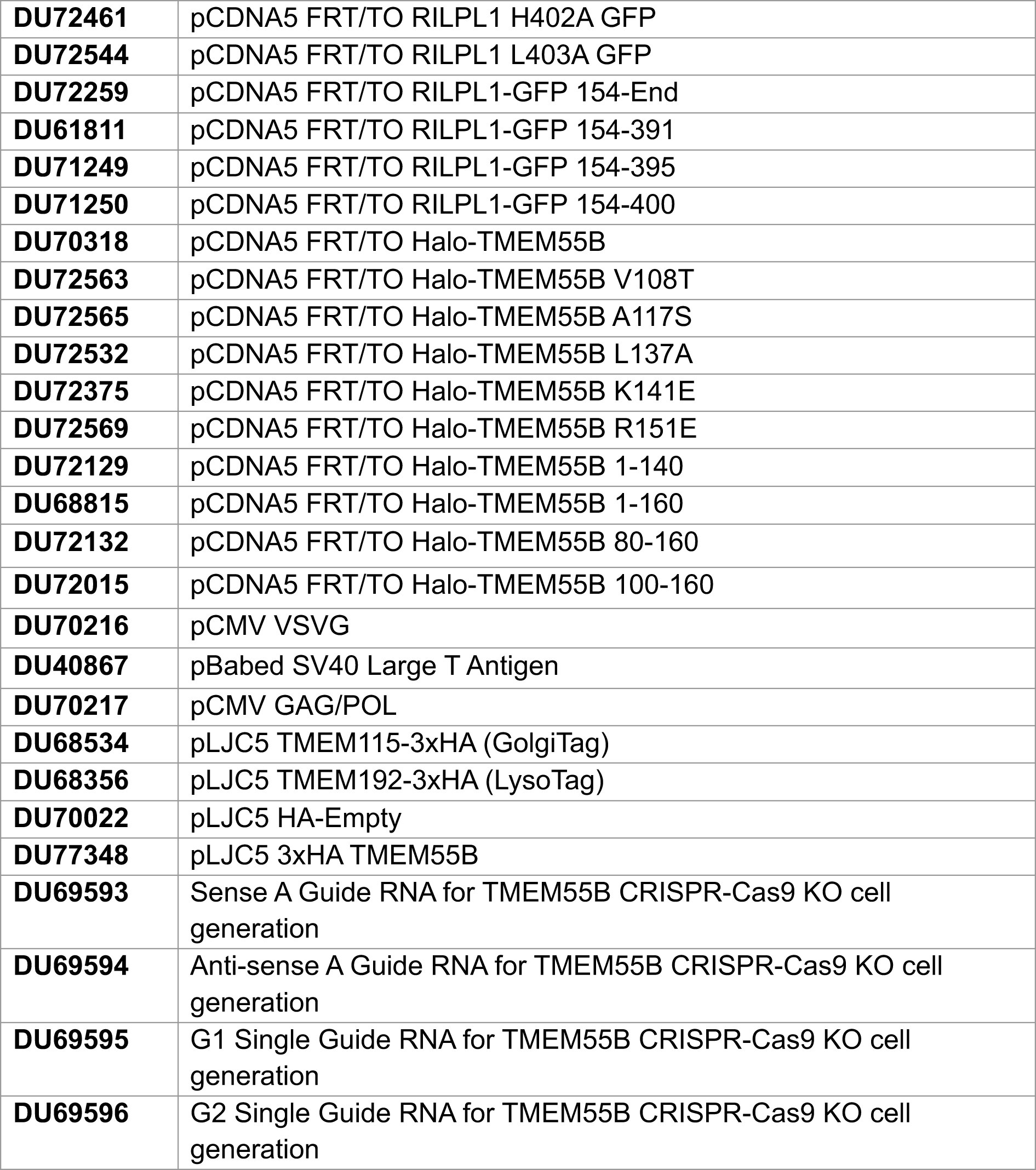

#### Primary Antibodies

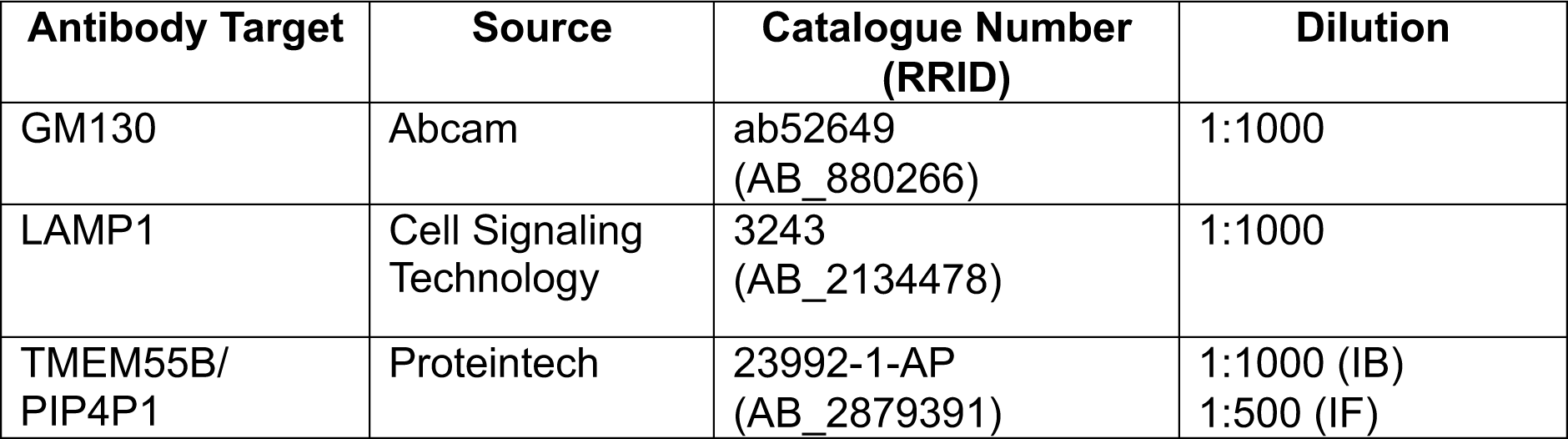
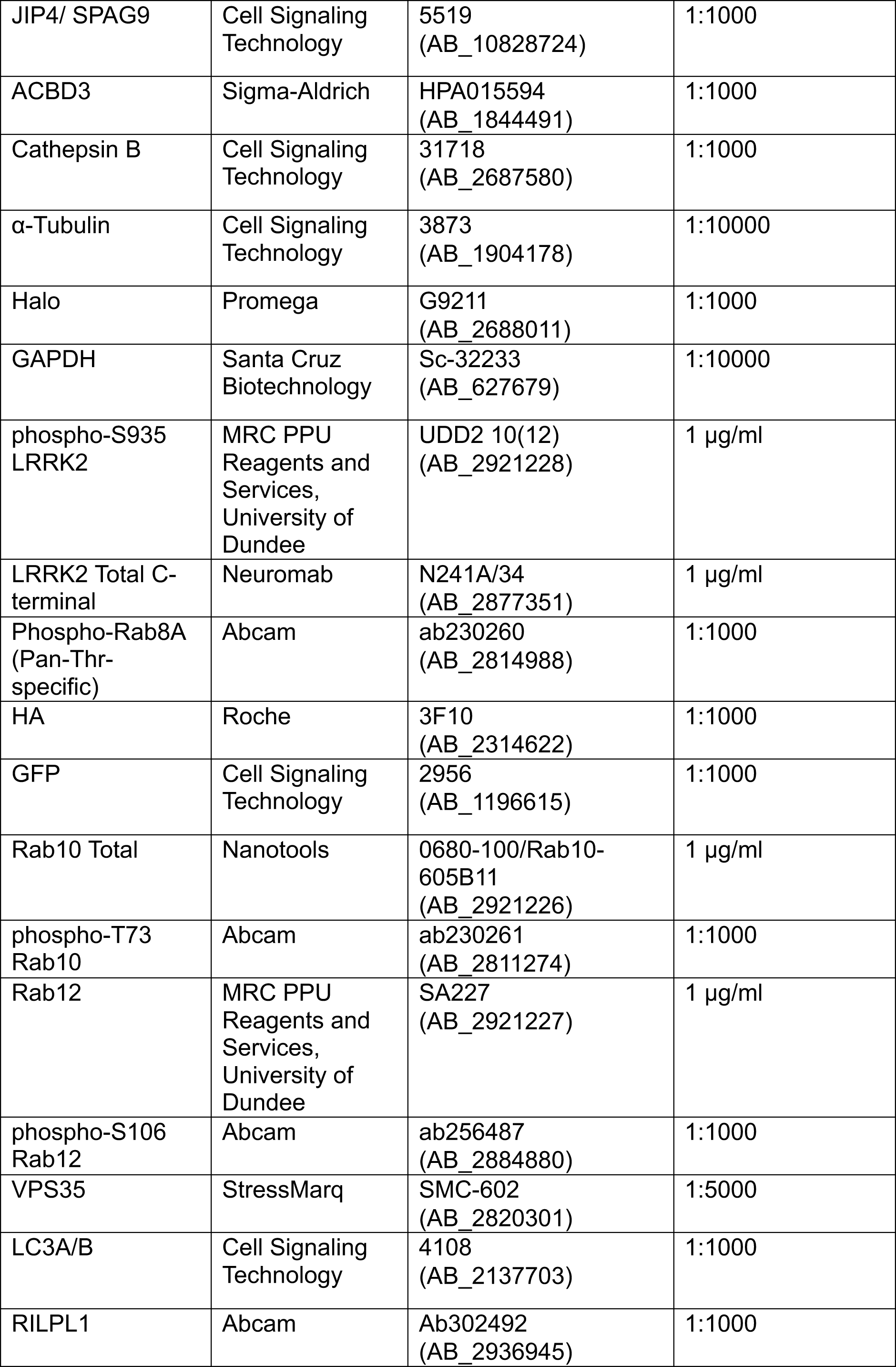
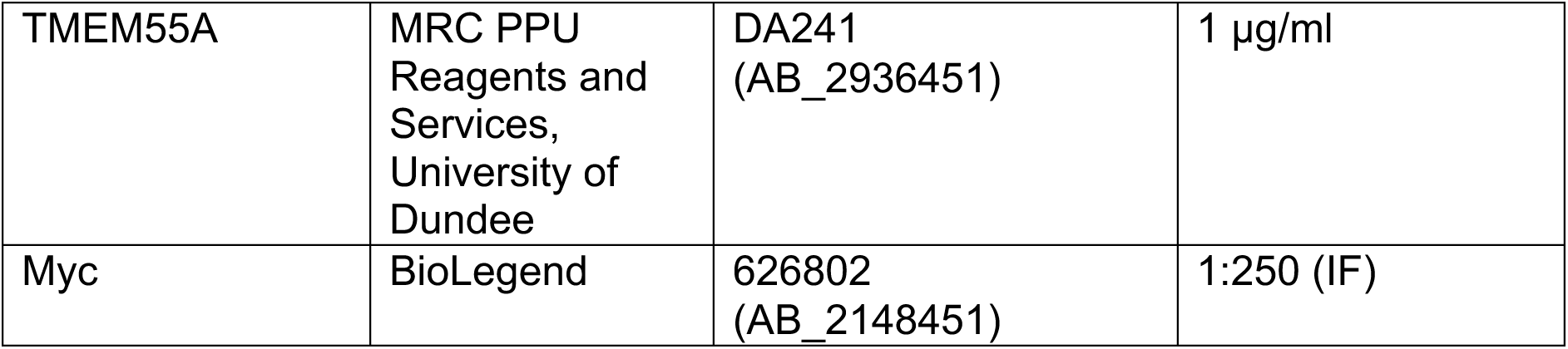

#### Secondary Antibodies

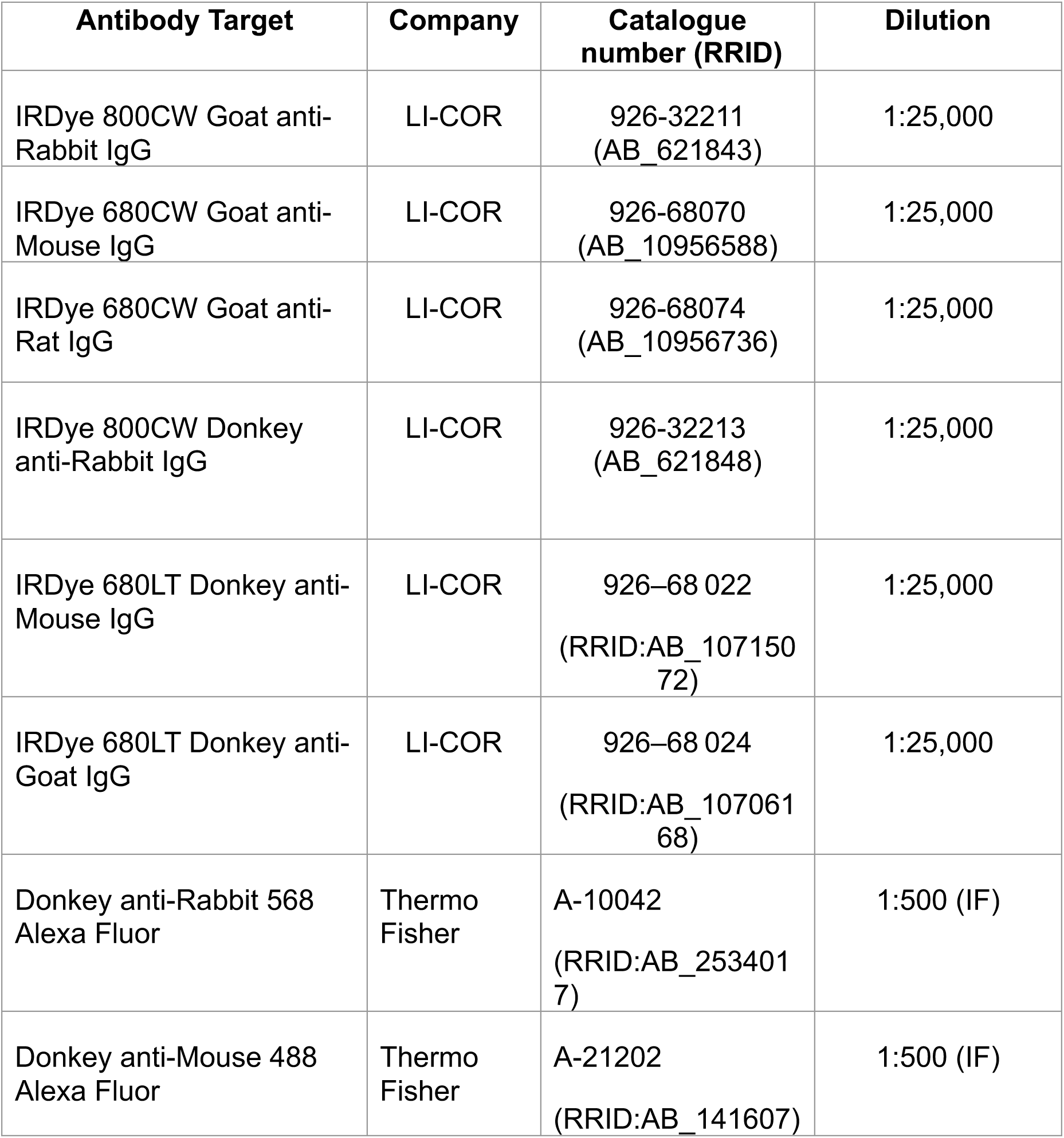

---

**Fig. S1.**
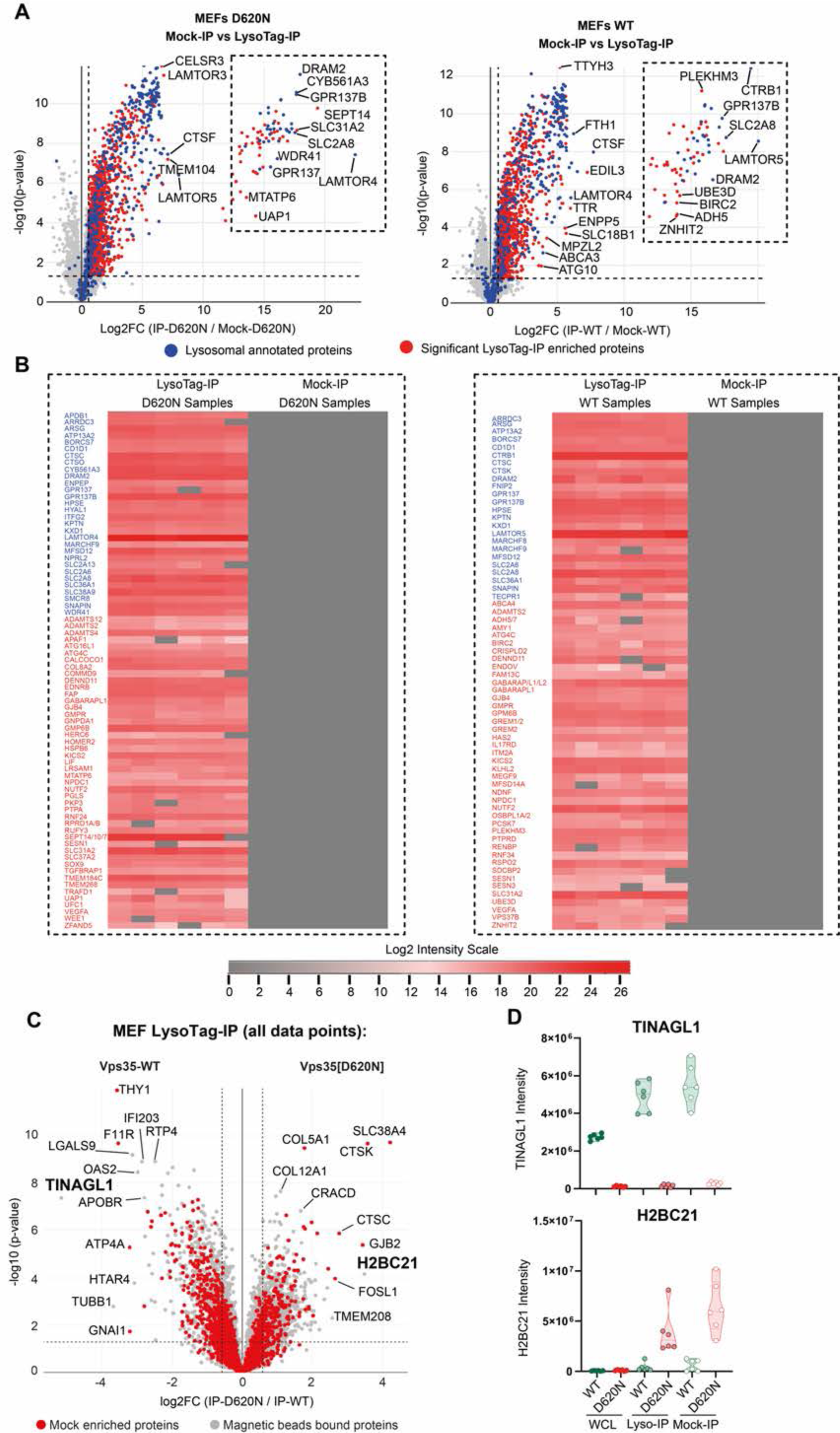
Further Mass Spectrometry analysis of wild type and D620N MEFs. (A) The volcano plot show the result of MEFs VPS35[D620N] not expressing LysoTag (Mock-IP) versus VPS35[D620N] LysoTag-IP (TMEM-192-3xHA) samples (left) (Curtain link: https://curtain.proteo.info/#/e7b85272-37e8-4785-9879-954f19c96784) and MEFs VPS35-WT not expressing LysoTag (Mock-IP) versus VPS35-WT LysoTag-IP samples (right) (Curtain link:https://curtain.proteo.info/#/4399d86e-67d2-40f6-af3c-6e237c54a589) (Table S1). The blue dots represent lysosomal proteins annotated in the GO terms database (GO:0005764), while the red dots represent significantly enriched proteins with fold change > 1.5 and p-value < 0.05. The black dotted line box indicates proteins confidently detected in the LysoTag-IP samples but not in the Mock-IP samples. (B) The log2 MS intensities of these proteins are shown. (C) The volcano plot (Curtain link: https://curtain.proteo.info/#/18c40d8a-f678-4d7e-976a-f70c85683ccc) shows the proteome changes of all detected proteins in MEFs VPS35-WT versus VPS35[D620N] LysoTag-IP experiments. The red dots represent mock-enriched proteins with fold change > 1.5 and p-value < 0.05, as determined in the experiments shown in Supplementary Figure 1A. The grey dots represent proteins bound to magnetic beads, which do not show significant enrichment. (D) The raw MS intensities of two non-specific proteins that associate with the magnetic beads (TINAGL1 and H2BC21) were shown for reference.

**Fig. S2.**
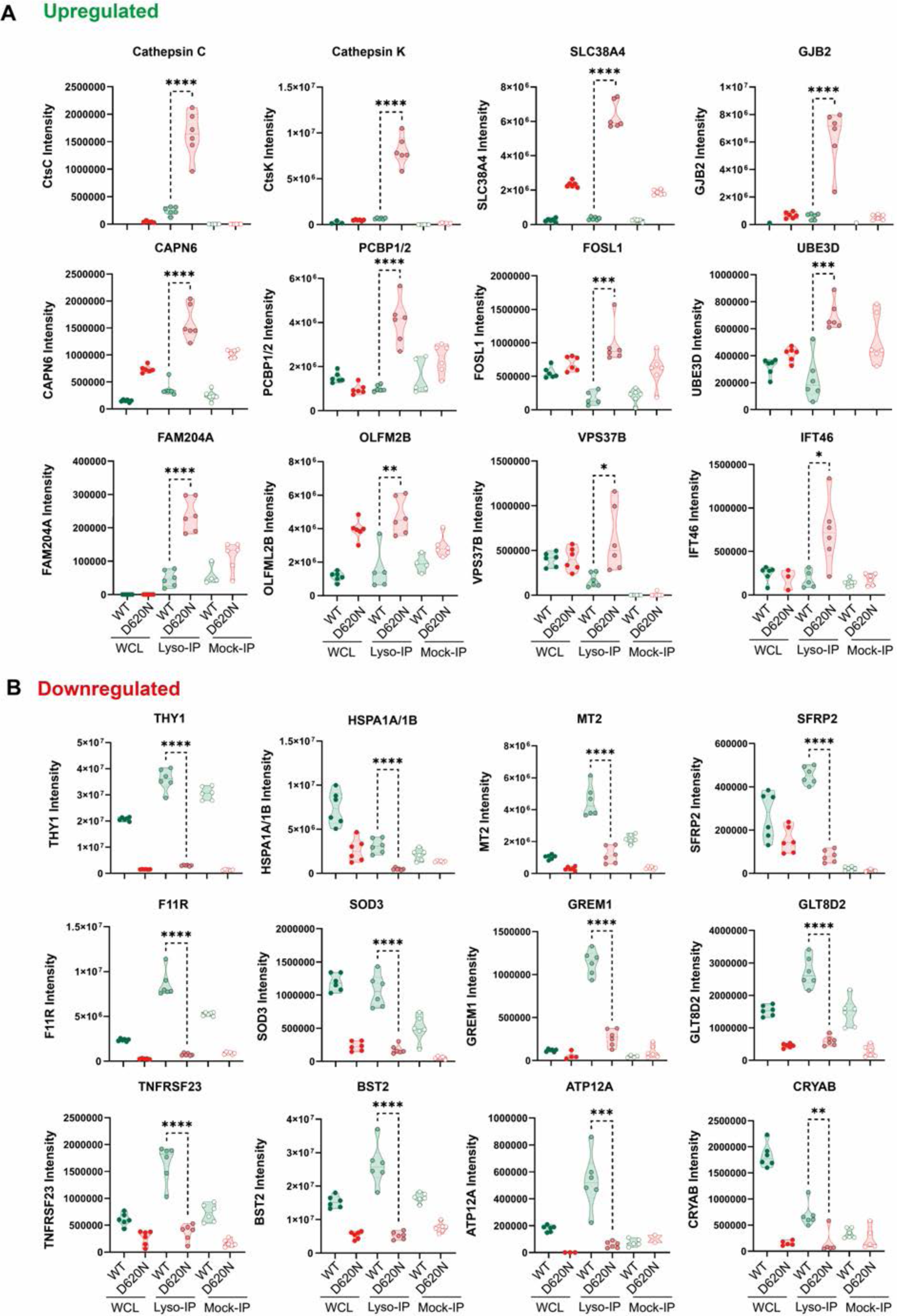
Relative expression of selected proteins in wild type and D620N whole cell lysates and lysosomes. Violin plots of the indicated proteins that are most (A) upregulated or (B) downregulated in the presence of VPS35[D620N] mutation are shown. Data were analyzed using two-tailed unpaired t-test (** p< 0.01, *** p< 0.001, **** p< 0.001).

**Fig. S3.**
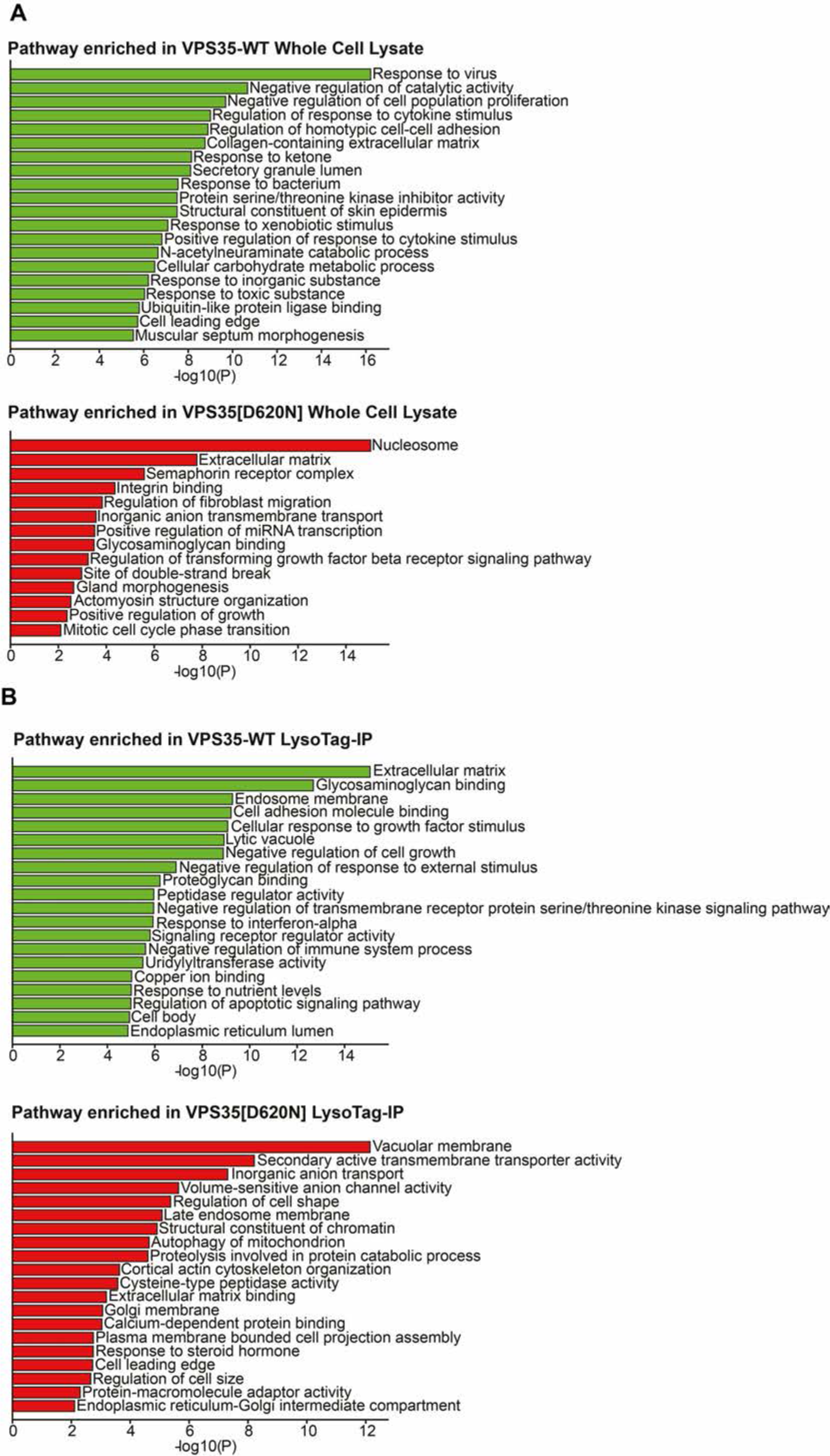
Gene Ontology analysis of proteins whose levels are most affected by the D620N mutations. The GO term pathway analysis of significant down-regulated (green bar) and up-regulated proteins (red bar) observed in (A) WCL (Fig. 1C) and (B) LysoTag-IP (Fig. 1D) experiments with fold change > 1.5 and p-value <0.05 using Metascape software (RRID:SCR_016620, version 5.3) with an enrichment of p-value < 0.01.

**Fig. S4.**
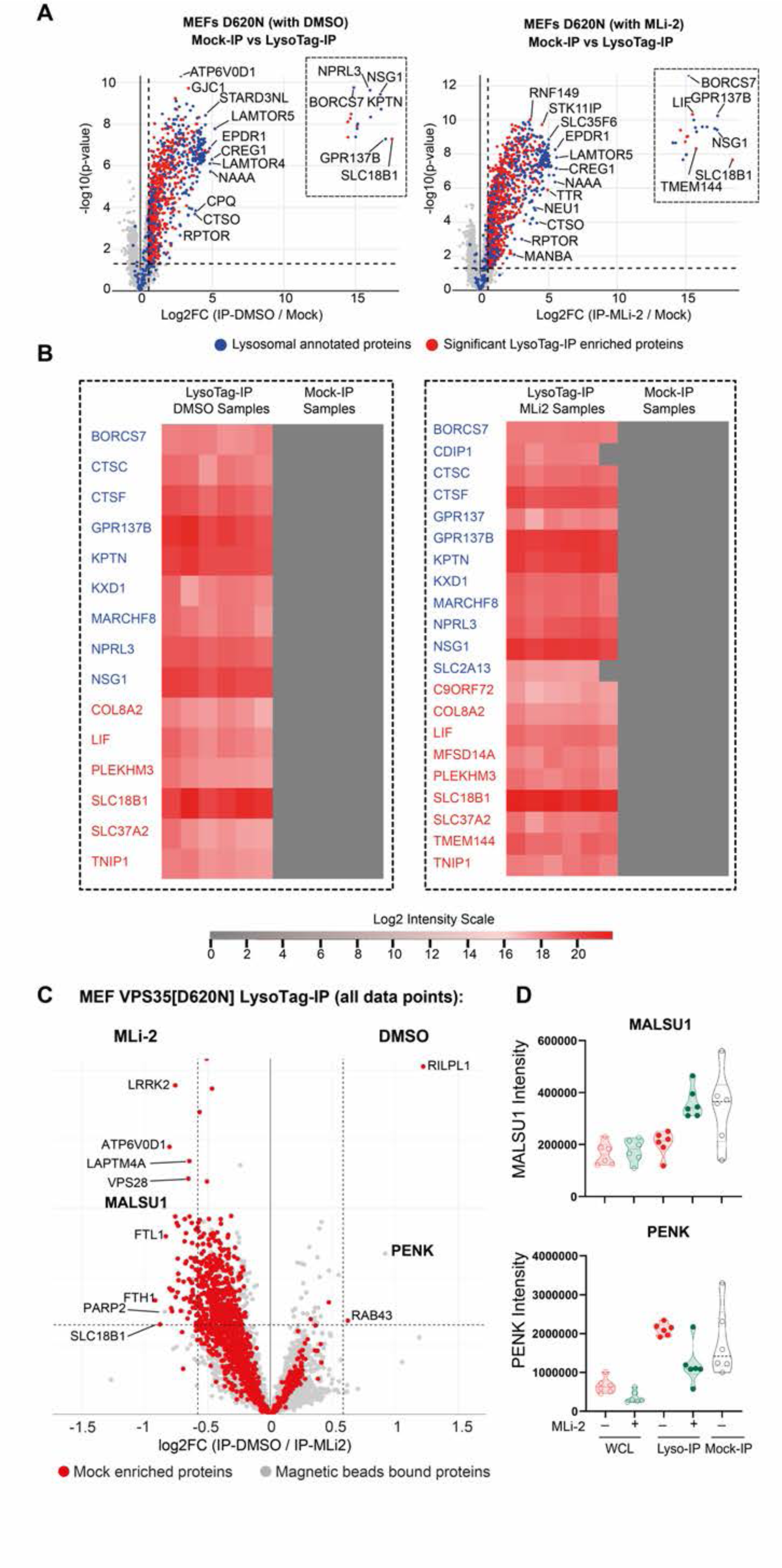
Further Mass Spectrometry analysis of D60N MEFs treated ± MLi-2. (A) The volcano plots show the result of MEFs VPS35[D620N] (no LysoTag) (Mock-IP) versus VPS35[D620N] (with DMSO) LysoTag-IP (TMEM192-3xHA) samples (left) (Curtain link: https://curtain.proteo.info/#/523b531b-eb5b-40a9-9f91-0aa7f09a002e) and MEFs VPS35[D620N] (no LysoTag) versus VPS35[D620N] (with MLi-2) LysoTag-IP samples (right) (Curtain link:https://curtain.proteo.info/#/100602aa-ca9d-4a5f-a821-4dc98b637570) (Table S1). The blue dots represent lysosomal proteins annotated in the GO terms database (GO:0005764), while the red dots represent significantly enriched proteins with fold change > 1.5 and p-value < 0.05. The black dotted line box indicates proteins confidently detected in the LysoTag-IP samples but not in the Mock-IP samples. The log2 MS intensities of these proteins are shown in Supplementary Figure (B). (C) The volcano plot (Curtain link:https://curtain.proteo.info/#/e10d5d9a-3214-4a5d-88cb-7ca1b1460e1c) shows the proteome changes of all detected proteins in MEFs VPS35[D620N] MLi-2 versus DMSO LysoTag-IP experiments. The red dots represent mock-enriched proteins with fold change > 1.5 and p-value < 0.05. The gray dots represent proteins bound to magnetic beads, which do not show significant enrichment. (D) The raw MS intensities of two non-specific proteins that associate with the magnetic beads (MALSU1 and PENK) were shown for reference.

**Fig. S5.**
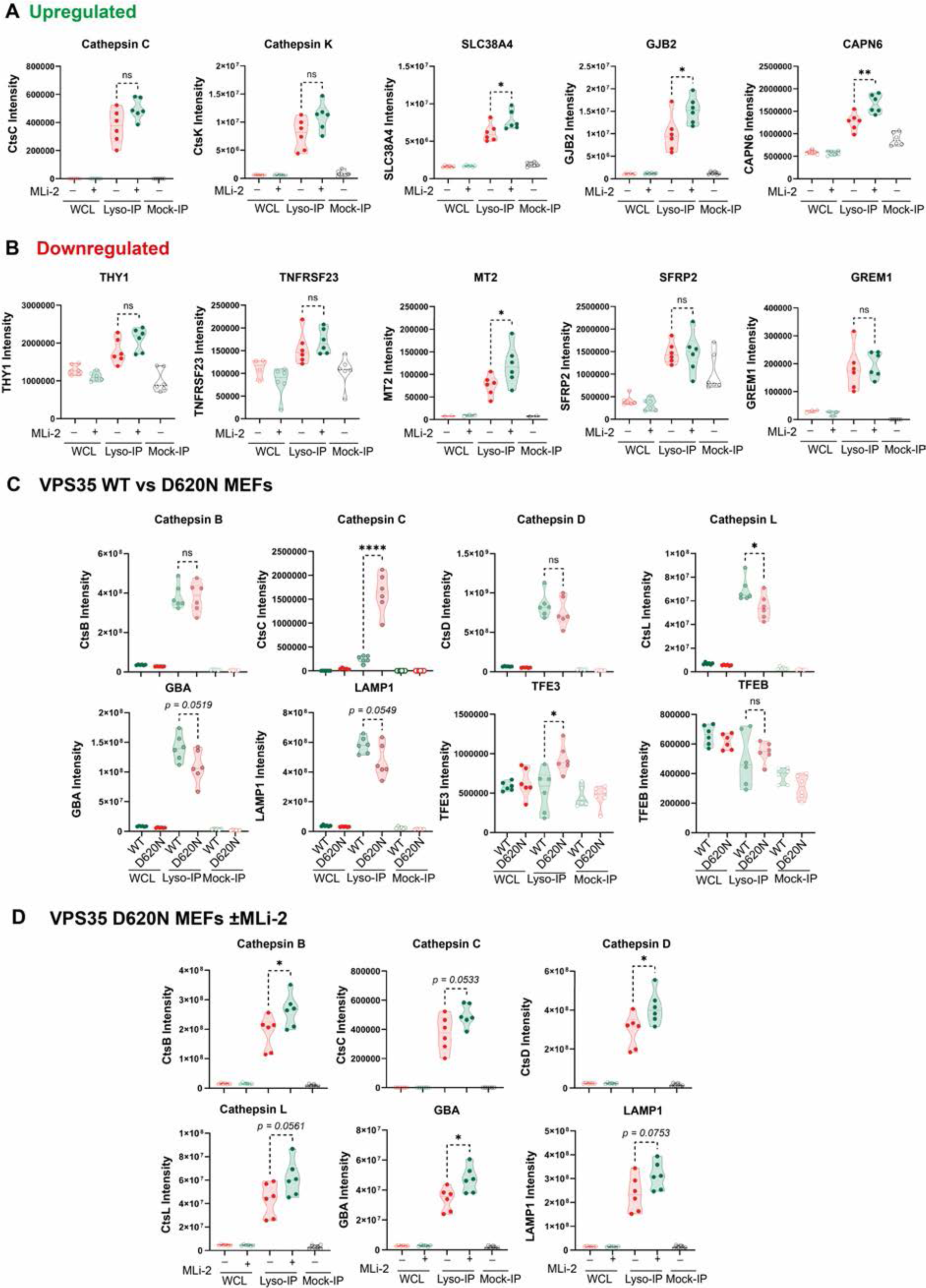
Relative levels of selected proteins in lysate and lysosomes of wild type and D620N MEFs and impact of MLi-2. (A to D) Violin plots of the indicated proteins from MS data presented in Fig. 1, A and C; and Fig. 2, B and D). Data were analyzed using two-tailed unpaired t-test (** p< 0.01, *** p< 0.001, **** p< 0.001).

**Fig.S6.**
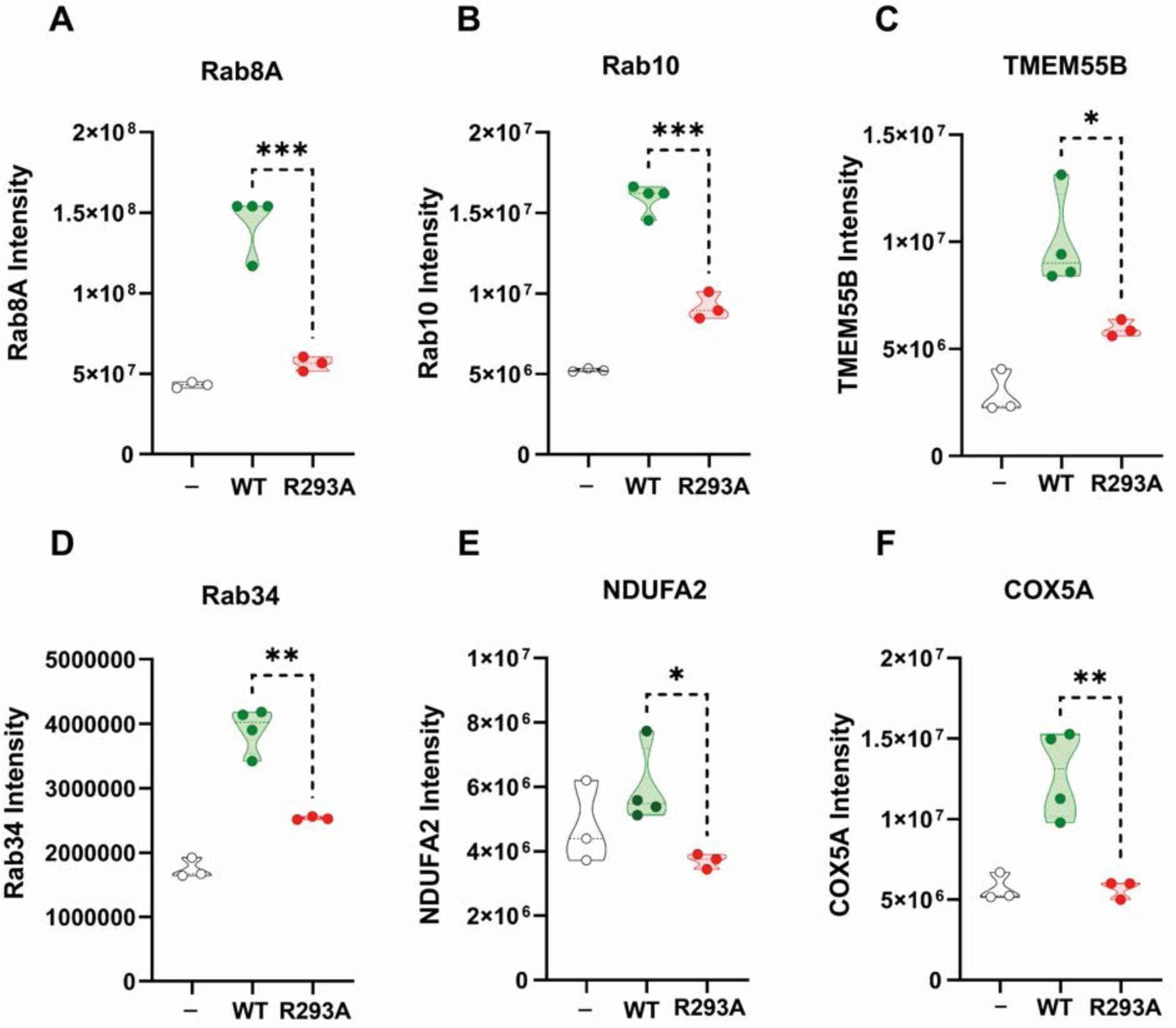
Relative levels of RILPL1 interacting proteins derived from Mass spectrometry data. (A to F) Violin plots of the indicated proteins from MS data presented in Fig. 4C. Data were analyzed using two-tailed unpaired t-test (*p< 0.05, ** p< 0.01, *** p< 0.001).

**Fig. S7.**
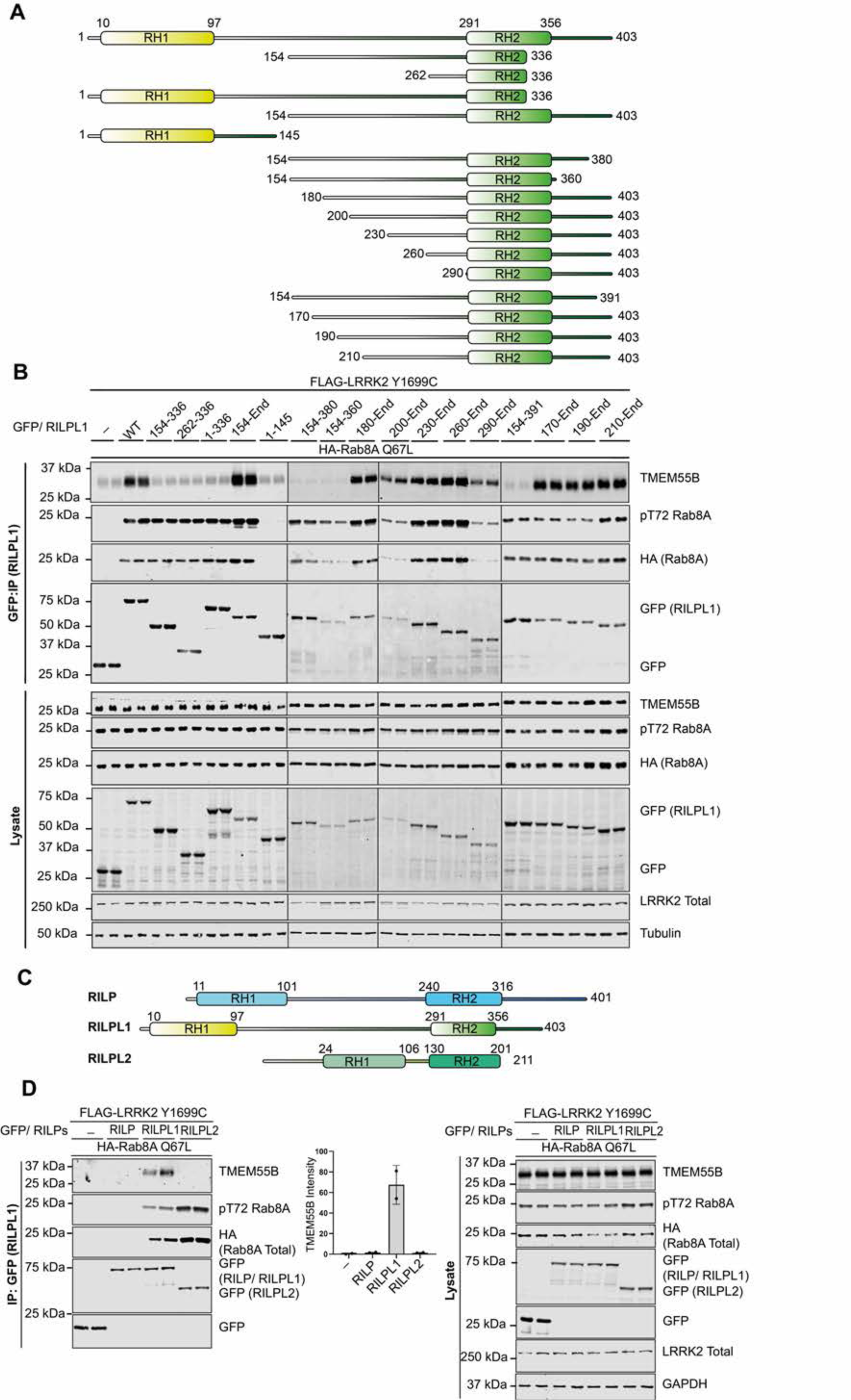
Further confirmation that RILPL1 interacts with TMEM55B but not RILP or RILPL2. (A) Domain structure of full length and truncated RILPL1 mutants. (B) HEK293 cells were transiently transfected with the indicated proteins and lysed 24h post transfection. GFP-RILPL1 immunoprecipitations (top panel) or cell extracts (lower panel) were subjected to quantitative immunoblot analysis using the LI-COR Odyssey CLx Western Blot imaging system and indicated antibodies. Quantitation of immunoblotting data (performed using ImageStudioLite software version 5.2.5, RRID:SCR_013715; http://www.licor.com/bio/products/software/image_studio_lite/) is shown as mean ± SEM. (C) Domain structure of full length RILP, RILPL1 and RILPL2. (D) As in (B).

**Fig. S8.**
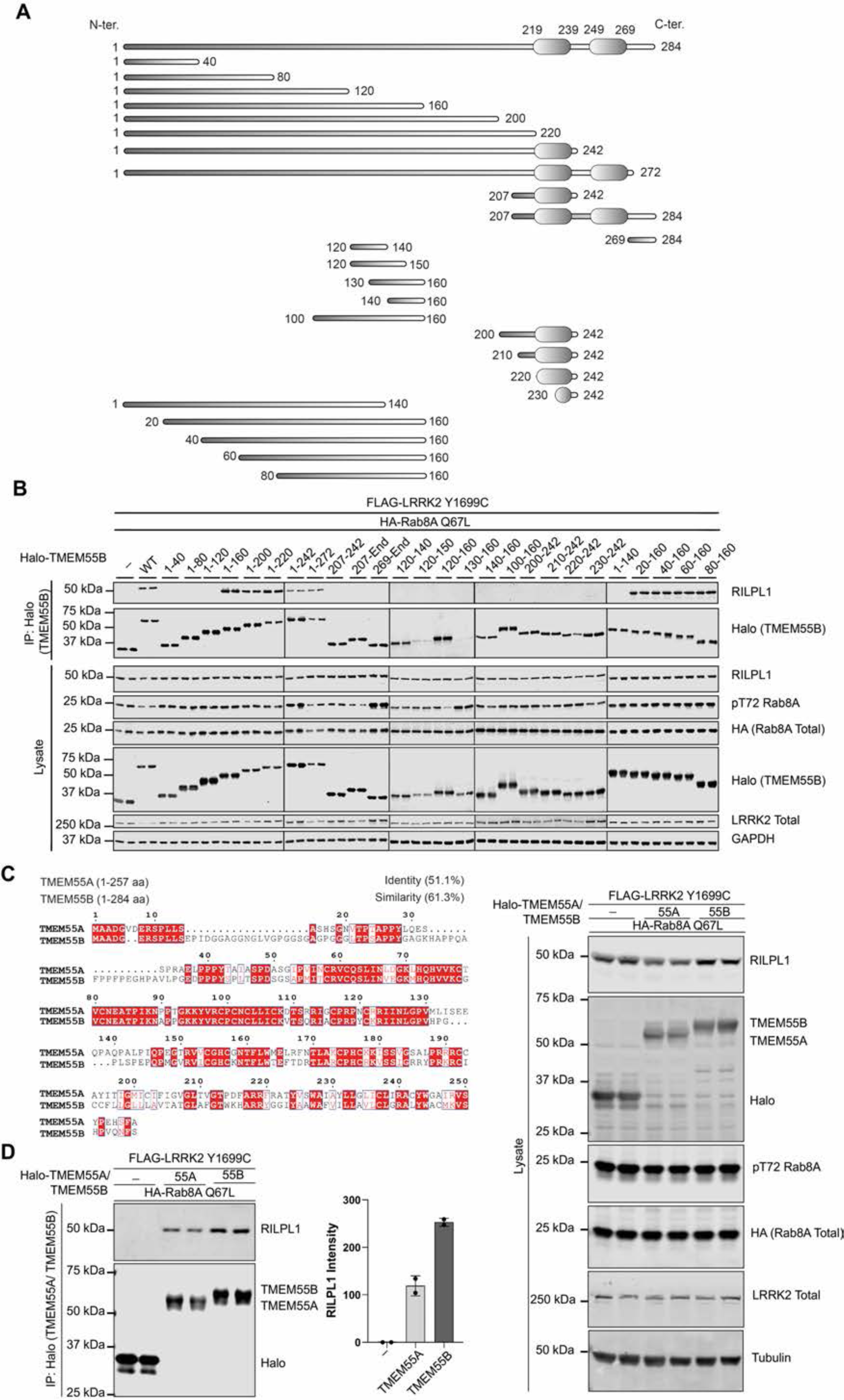
Conformation that the TMEM55B Conserved domain binds RILPL1 and that TMEM55A also binds RILPL1. (A) Domain structure of full length and truncated mutants of TMEM55B used in this study. (B) HEK293 cells were transiently transfected with the indicated proteins and lysed 24h post transfection. Halo-TMEM55B immunoprecipitations (top panel) or cell extracts (lower panel) were subjected to quantitative immunoblot analysis using the LI-COR Odyssey CLx Western Blot imaging system and indicated antibodies. (C) Multiple sequence alignment of TMEM55A and TMEM55B using Clustal Omega (https://www.ebi.ac.uk/Tools/msa/clustalo/) and ESPript (https://espript.ibcp.fr/ESPript/cgi-bin/ESPript.cgi) (*65*) (D) As in (B).

**Fig. S9.**
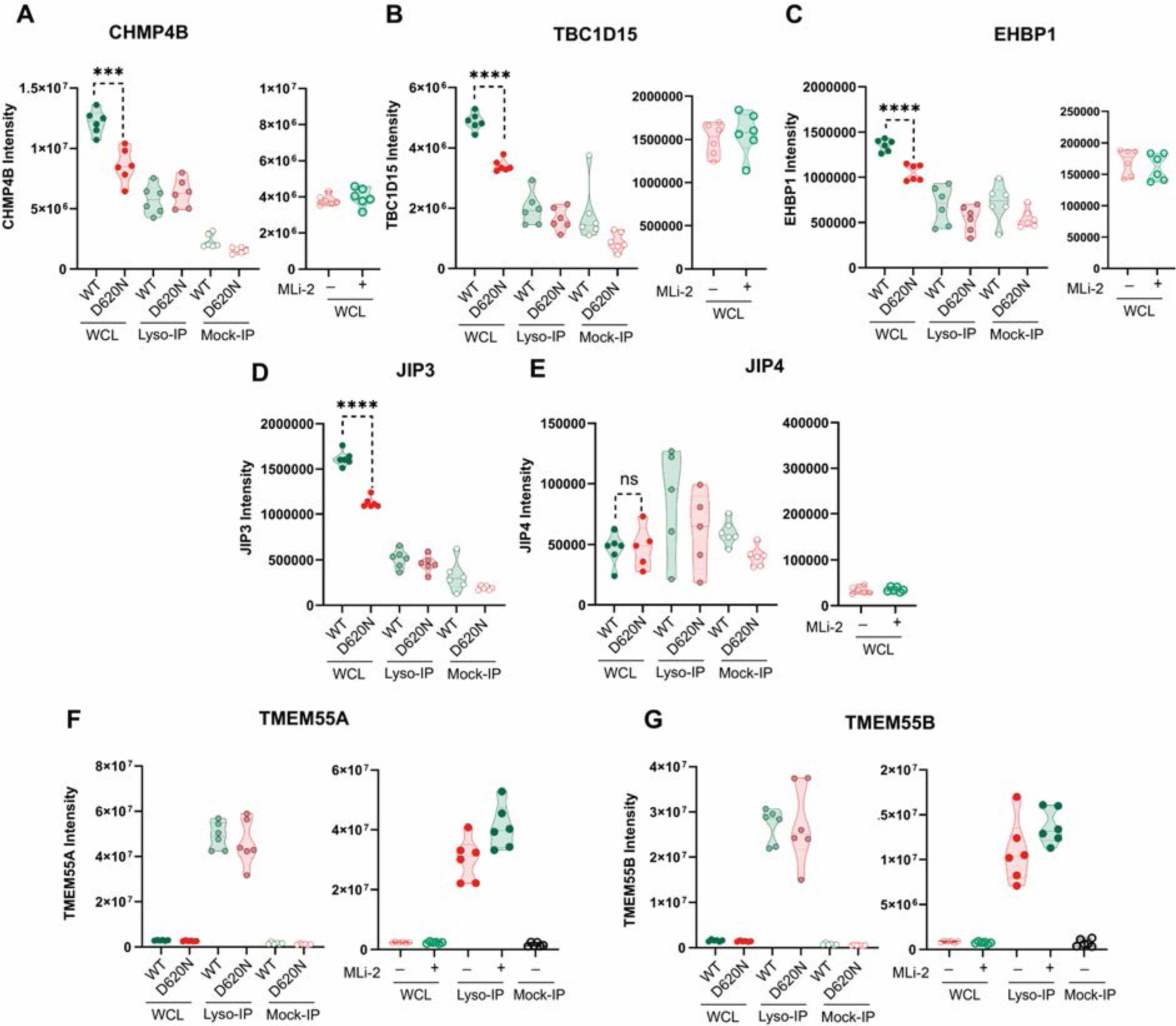
Relative levels of proteins in D620N Lysates and lysosomes that have been reported to be recruited to the lysosome in LLOMe treated cells. (A to G) Violin plots of the indicated proteins from MS data presented in Figure 1 (data on left panels) and Figure 2 (data on right panels). Data were analyzed using two-tailed unpaired *t*-test (*** p< 0.001, **** p< 0.001).

## Legends for Tables S1 to S3 that are Excel sheets submitted separately as files

Table S1. Search result of DIA MS data: VPS35 WT vs D620N WCL and LysoTag IP

Table S2. Search result of DIA MS data: VPS35 D620N ± MLi-2 WCL and LysoTag IP

Table S3. Search result of TMT MS data: RILPL1 WT vs R293A GFP I

## References

1. D. R. Alessi, E. Sammler, LRRK2 kinase in Parkinson’s disease. Science 360, 36–37 (2018).

2. E. Tolosa, M. Vila, C. Klein, O. Rascol, LRRK2 in Parkinson disease: challenges of clinical trials. Nat Rev Neurol 16, 97–107 (2020).

3. M. K. Herrick, M. G. Tansey, Is LRRK2 the missing link between inflammatory bowel disease and Parkinson’s disease? NPJ Parkinsons Dis 7, 26 (2021).

4. K. Y. Hui, H. Fernandez-Hernandez, J. Hu, A. Schaffner, N. Pankratz, N. Y. Hsu, L. S. Chuang, S. Carmi, N. Villaverde, X. Li, M. Rivas, A. P. Levine, X. Bao, P. R. Labrias, T. Haritunians, D. Ruane, K. Gettler, E. Chen, D. Li, E. R. Schiff, N. Pontikos, N. Barzilai, S. R. Brant, S. Bressman, A. S. Cheifetz, L. N. Clark, M. J. Daly, R. J. Desnick, R. H. Duerr, S. Katz, T. Lencz, R. H. Myers, H. Ostrer, L. Ozelius, H. Payami, Y. Peter, J. D. Rioux, A. W. Segal, W. K. Scott, M. S. Silverberg, J. M. Vance, I. Ubarretxena-Belandia, T. Foroud, G. Atzmon, I. Pe’er, Y. Ioannou, D. P. B. McGovern, Z. Yue, E. E. Schadt, J. H. Cho, I. Peter, Functional variants in the LRRK2 gene confer shared effects on risk for Crohn’s disease and Parkinson’s disease. Sci Transl Med 10, (2018).

5. M. Taylor, D. R. Alessi, Advances in elucidating the function of leucine-rich repeat protein kinase-2 in normal cells and Parkinson’s disease. Curr Opin Cell Biol 63, 102–113 (2020).

6. M. Steger, F. Diez, H. S. Dhekne, P. Lis, R. S. Nirujogi, O. Karayel, F. Tonelli, T. N. Martinez, E. Lorentzen, S. R. Pfeffer, D. R. Alessi, M. Mann, Systematic proteomic analysis of LRRK2-mediated Rab GTPase phosphorylation establishes a connection to ciliogenesis. Elife 6, e31012 (2017).

7. M. Steger, F. Tonelli, G. Ito, P. Davies, M. Trost, M. Vetter, S. Wachter, E. Lorentzen, G. Duddy, S. Wilson, M. A. Baptista, B. K. Fiske, M. J. Fell, J. A. Morrow, A. D. Reith, D. R. Alessi, M. Mann, Phosphoproteomics reveals that Parkinson’s disease kinase LRRK2 regulates a subset of Rab GTPases. Elife 5, e12813 (2016).

8. S. R. Pfeffer, Rab GTPases: master regulators that establish the secretory and endocytic pathways. Mol Biol Cell 28, 712–715 (2017).

9. K. Berndsen, P. Lis, W. M. Yeshaw, P. S. Wawro, R. S. Nirujogi, M. Wightman, T. Macartney, M. Dorward, A. Knebel, F. Tonelli, S. R. Pfeffer, D. R. Alessi, PPM1H phosphatase counteracts LRRK2 signaling by selectively dephosphorylating Rab proteins. eLife 8, e50416 (2019).

10. D. Waschbüsch, K. Berndsen, A. Knebel, D. R. Alessi, A. R. Khan, Structural basis for the specificity of PPM1H phosphatase for Rab GTPases. bioRxiv, 2021.2002.2017.431620 (2021).

11. D. Waschbüsch, E. Purlyte, P. Pal, E. McGrath, D. R. Alessi, A. R. Khan, Structural Basis for Rab8a Recruitment of RILPL2 via LRRK2 Phosphorylation of Switch 2. Structure 28, 406–417.e406 (2020).

12. L. Bonet-Ponce, A. Beilina, C. D. Williamson, E. Lindberg, J. H. Kluss, S. Saez-Atienzar, N. Landeck, R. Kumaran, A. Mamais, C. K. E. Bleck, Y. Li, M. R. Cookson, LRRK2 mediates tubulation and vesicle sorting from lysosomes. Sci Adv 6, (2020).

13. Y. Sobu, P. S. Wawro, H. S. Dhekne, S. R. Pfeffer, Pathogenic LRRK2 regulates ciliation probability upstream of Tau Tubulin kinase 2. bioRxiv, 2020.2004.2007.029983 (2020).

14. H. S. Dhekne, I. Yanatori, R. C. Gomez, F. Tonelli, F. Diez, B. Schule, M. Steger, D. R. Alessi, S. R. Pfeffer, A pathway for Parkinson’s Disease LRRK2 kinase to block primary cilia and Sonic hedgehog signaling in the brain. Elife 7, e40202 (2018).

15. E. Purlyte, H. S. Dhekne, A. R. Sarhan, R. Gomez, P. Lis, M. Wightman, T. N. Martinez, F. Tonelli, S. R. Pfeffer, D. R. Alessi, Rab29 activation of the Parkinson’s disease-associated LRRK2 kinase. EMBO J 37, 1–18 (2018).

16. Z. Liu, N. Bryant, R. Kumaran, A. Beilina, A. Abeliovich, M. R. Cookson, A. B. West, LRRK2 phosphorylates membrane-bound Rabs and is activated by GTP-bound Rab7L1 to promote recruitment to the trans-Golgi network. Hum Mol Genet 27, 385–395 (2018).

17. E. McGrath, D. Waschbüsch, B. M. Baker, A. R. Khan, LRRK2 binds to the Rab32 subfamily in a GTP-dependent manner via its armadillo domain. Small GTPases, 1–14 (2019).

18. E. G. Vides, A. Adhikari, C. Y. Chiang, P. Lis, E. Purlyte, C. Limouse, J. L. Shumate, E. Spinola-Lasso, H. S. Dhekne, D. R. Alessi, S. R. Pfeffer, A feed-forward pathway drives LRRK2 kinase membrane recruitment and activation. Elife 11, (2022).

19. H. Zhu, F. Tonelli, D. R. Alessi, J. Sun, Structural basis of human LRRK2 membrane recruitment and activation. bioRxiv, 2022.2004.2026.489605 (2022).

20. H. S. Dhekne, F. Tonelli, W. M. Yeshaw, C. Y. Chiang, C. Limouse, E. Jaimon, E. Purlyte, D. R. Alessi, S. R. Pfeffer, Genome-wide screen reveals Rab12 GTPase as a critical activator of pathogenic LRRK2 kinase. bioRxiv, 2023.2002.2017.529028 (2023).

21. V. V. Bondar, X. Wang, O. B. Davis, M. T. Maloney, M. Agam, M. Y. Chin, A. C.-N. Ho, D. Joy, J. W. Lewcock, G. D. Paolo, R. G. Thorne, Z. K. Sweeney, A. G. Henry, Rab12 regulates LRRK2 activity by promoting its localization to lysosomes. bioRxiv, 2023.2002.2021.529466 (2023).

22. G. K. Tofaris, Lysosome-dependent pathways as a unifying theme in Parkinson’s disease. Mov Disord 27, 1364–1369 (2012).

23. A. Navarro-Romero, M. Montpeyo, M. Martinez-Vicente, The Emerging Role of the Lysosome in Parkinson’s Disease. Cells 9, (2020).

24. E. Sidransky, M. A. Nalls, J. O. Aasly, J. Aharon-Peretz, G. Annesi, E. R. Barbosa, A. Bar-Shira, D. Berg, J. Bras, A. Brice, C. M. Chen, L. N. Clark, C. Condroyer, E. V. De Marco, A. Durr, M. J. Eblan, S. Fahn, M. J. Farrer, H. C. Fung, Z. Gan-Or, T. Gasser, R. Gershoni-Baruch, N. Giladi, A. Griffith, T. Gurevich, C. Januario, P. Kropp, A. E. Lang, G. J. Lee-Chen, S. Lesage, K. Marder, I. F. Mata, A. Mirelman, J. Mitsui, I. Mizuta, G. Nicoletti, C. Oliveira, R. Ottman, A. Orr-Urtreger, L. V. Pereira, A. Quattrone, E. Rogaeva, A. Rolfs, H. Rosenbaum, R. Rozenberg, A. Samii, T. Samaddar, C. Schulte, M. Sharma, A. Singleton, M. Spitz, E. K. Tan, N. Tayebi, T. Toda, A. R. Troiano, S. Tsuji, M. Wittstock, T. G. Wolfsberg, Y. R. Wu, C. P. Zabetian, Y. Zhao, S. G. Ziegler, Multicenter analysis of glucocerebrosidase mutations in Parkinson’s disease. N Engl J Med 361, 1651–1661 (2009).

25. S. van Veen, S. Martin, C. Van den Haute, V. Benoy, J. Lyons, R. Vanhoutte, J. P. Kahler, J. P. Decuypere, G. Gelders, E. Lambie, J. Zielich, J. V. Swinnen, W. Annaert, P. Agostinis, B. Ghesquiere, S. Verhelst, V. Baekelandt, J. Eggermont, P. Vangheluwe, ATP13A2 deficiency disrupts lysosomal polyamine export. Nature 578, 419–424 (2020).

26. M. Hu, P. Li, C. Wang, X. Feng, Q. Geng, W. Chen, M. Marthi, W. Zhang, C. Gao, W. Reid, J. Swanson, W. Du, R. I. Hume, H. Xu, Parkinson’s disease-risk protein TMEM175 is a proton-activated proton channel in lysosomes. Cell 185, 2292–2308 e2220 (2022).

27. M. Madureira, N. Connor-Robson, R. Wade-Martins, “LRRK2: Autophagy and Lysosomal Activity”. Front Neurosci 14, 498 (2020).

28. C. A. Boecker, J. Goldsmith, D. Dou, G. G. Cajka, E. L. F. Holzbaur, Increased LRRK2 kinase activity alters neuronal autophagy by disrupting the axonal transport of autophagosomes. Curr Biol 31, 2140–2154 e2146 (2021).

29. F. Singh, A. R. Prescott, P. Rosewell, G. Ball, A. D. Reith, I. G. Ganley, Pharmacological rescue of impaired mitophagy in Parkinson’s disease-related LRRK2 G2019S knock-in mice. Elife 10, (2021).

30. S. Herbst, P. Campbell, J. Harvey, E. M. Bernard, V. Papayannopoulos, N. W. Wood, H. R. Morris, M. G. Gutierrez, LRRK2 activation controls the repair of damaged endomembranes in macrophages. Embo j, e104494 (2020).

31. A. F. Kalogeropulou, J. B. Freemantle, P. Lis, E. G. Vides, N. K. Polinski, D. R. Alessi, Endogenous Rab29 does not impact basal or stimulated LRRK2 pathway activity. Biochem J 477, 4397–4423 (2020).

32. N. Yadavalli, S. M. Ferguson, LRRK2 Suppresses Lysosome Degradative Activity in Macrophages and Microglia via Transcription Factor E3 Inhibition. bioRxiv, 2022.2012.2017.520834 (2022).

33. R. N. Alcalay, F. Hsieh, E. Tengstrand, S. Padmanabhan, M. Baptista, C. Kehoe, S. Narayan, A. K. Boehme, K. Merchant, Higher Urine bis(Monoacylglycerol)Phosphate Levels in LRRK2 G2019S Mutation Carriers: Implications for Therapeutic Development. Mov Disord 35, 134–141 (2020).

34. M. T. Maloney, X. Wang, R. Ghosh, S. V. Andrews, R. Maciuca, S. T. Masoud, R. M. Caprioli, J. Chen, C.-L. Chiu, S. S. Davis, A. C.-N. Ho, H. N. Nguyen, N. E. Propson, M. L. Reyzer, O. B. Davis, M. C. Deen, S. Zhu, G. Di Paolo, D. J. Vocadlo, A. A. Estrada, J. de Vicente, J. W. Lewcock, A. Arguello, J. H. Suh, S. Huntwork-Rodriguez, A. G. Henry, LRRK2 Kinase Activity Regulates Parkinson’s Disease-Relevant Lipids at the Lysosome. bioRxiv, 2022.2012.2019.521070 (2022).

35. A. Zimprich, A. Benet-Pages, W. Struhal, E. Graf, S. H. Eck, M. N. Offman, D. Haubenberger, S. Spielberger, E. C. Schulte, P. Lichtner, S. C. Rossle, N. Klopp, E. Wolf, K. Seppi, W. Pirker, S. Presslauer, B. Mollenhauer, R. Katzenschlager, T. Foki, C. Hotzy, E. Reinthaler, A. Harutyunyan, R. Kralovics, A. Peters, F. Zimprich, T. Brucke, W. Poewe, E. Auff, C. Trenkwalder, B. Rost, G. Ransmayr, J. Winkelmann, T. Meitinger, T. M. Strom, A mutation in VPS35, encoding a subunit of the retromer complex, causes late-onset Parkinson disease. Am J Hum Genet 89, 168–175 (2011).

36. C. Vilarino-Guell, C. Wider, O. A. Ross, J. C. Dachsel, J. M. Kachergus, S. J. Lincoln, A. I. Soto-Ortolaza, S. A. Cobb, G. J. Wilhoite, J. A. Bacon, B. Behrouz, H. L. Melrose, E. Hentati, A. Puschmann, D. M. Evans, E. Conibear, W. W. Wasserman, J. O. Aasly, P. R. Burkhard, R. Djaldetti, J. Ghika, F. Hentati, A. Krygowska-Wajs, T. Lynch, E. Melamed, A. Rajput, A. H. Rajput, A. Solida, R. M. Wu, R. J. Uitti, Z. K. Wszolek, F. Vingerhoets, M. J. Farrer, VPS35 mutations in Parkinson disease. Am J Hum Genet 89, 162–167 (2011).

37. S. A. Small, K. Kent, A. Pierce, C. Leung, M. S. Kang, H. Okada, L. Honig, J. P. Vonsattel, T. W. Kim, Model-guided microarray implicates the retromer complex in Alzheimer’s disease. Ann Neurol 58, 909–919 (2005).

38. R. Mir, F. Tonelli, P. Lis, T. Macartney, N. K. Polinski, T. N. Martinez, M.-Y. Chou, A. J. M. Howden, T. König, C. Hotzy, I. Milenkovic, T. Brücke, A. Zimprich, E. Sammler, D. R. Alessi, The Parkinson’s disease VPS35[D620N] mutation enhances LRRK2-mediated Rab protein phosphorylation in mouse and human. The Biochemical journal 475, 1861–1883 (2018).

39. R. S. Nirujogi, F. Tonelli, M. Taylor, P. Lis, A. Zimprich, E. Sammler, D. R. Alessi, Development of a multiplexed targeted mass spectrometry assay for LRRK2-phosphorylated Rabs and Ser910/Ser935 biomarker sites. Biochem J 478, 299–326 (2021).

40. T. S. Gomez, D. D. Billadeau, A FAM21-containing WASH complex regulates retromer-dependent sorting. Dev Cell 17, 699–711 (2009).

41. Y. Cui, Z. Yang, R. D. Teasdale, The functional roles of retromer in Parkinson’s disease. FEBS Lett, (2017).

42. M. Abu-Remaileh, G. A. Wyant, C. Kim, N. N. Laqtom, M. Abbasi, S. H. Chan, E. Freinkman, D. M. Sabatini, Lysosomal metabolomics reveals V-ATPase- and mTOR-dependent regulation of amino acid efflux from lysosomes. Science (New York, N.Y.) 358, 807–813 (2017).

43. V. Demichev, C. B. Messner, S. I. Vernardis, K. S. Lilley, M. Ralser, DIA-NN: neural networks and interference correction enable deep proteome coverage in high throughput. Nat Methods 17, 41–44 (2020).

44. Y. Zhou, B. Zhou, L. Pache, M. Chang, A. H. Khodabakhshi, O. Tanaseichuk, C. Benner, S. K. Chanda, Metascape provides a biologist-oriented resource for the analysis of systems-level datasets. Nat Commun 10, 1523 (2019).

45. R. Fasimoye, W. Dong, R. S. Nirujogi, E. S. Rawat, M. Iguchi, K. Nyame, T. K. Phung, E. Bagnoli, A. R. Prescott, D. R. Alessi, M. Abu-Remaileh, Golgi-IP, a tool for multimodal analysis of Golgi molecular content. Proc Natl Acad Sci U S A 120, e2219953120 (2023).

46. M. J. Fell, C. Mirescu, K. Basu, B. Cheewatrakoolpong, D. E. DeMong, J. M. Ellis, L. A. Hyde, Y. Lin, C. G. Markgraf, H. Mei, M. Miller, F. M. Poulet, J. D. Scott, M. D. Smith, Z. Yin, X. Zhou, E. M. Parker, M. E. Kennedy, J. A. Morrow, MLi-2, a Potent, Selective, and Centrally Active Compound for Exploring the Therapeutic Potential and Safety of LRRK2 Kinase Inhibition. J Pharmacol Exp Ther 355, 397–409 (2015).

47. G. Ito, K. Katsemonova, F. Tonelli, P. Lis, M. A. Baptista, N. Shpiro, G. Duddy, S. Wilson, P. W. Ho, S. L. Ho, A. D. Reith, D. R. Alessi, Phos-tag analysis of Rab10 phosphorylation by LRRK2: a powerful assay for assessing kinase function and inhibitors. Biochem J 473, 2671–2685 (2016).

48. R. Willett, J. A. Martina, J. P. Zewe, R. Wills, G. R. V. Hammond, R. Puertollano, TFEB regulates lysosomal positioning by modulating TMEM55B expression and JIP4 recruitment to lysosomes. Nat Commun 8, 1580 (2017).

49. I. J. McGough, F. Steinberg, D. Jia, P. A. Barbuti, K. J. McMillan, K. J. Heesom, A. L. Whone, M. A. Caldwell, D. D. Billadeau, M. K. Rosen, P. J. Cullen, Retromer binding to FAM21 and the WASH complex is perturbed by the Parkinson disease-linked VPS35(D620N) mutation. Curr Biol 24, 1670–1676 (2014).

50. E. Zavodszky, M. N. Seaman, K. Moreau, M. Jimenez-Sanchez, S. Y. Breusegem, M. E. Harbour, D. C. Rubinsztein, Mutation in VPS35 associated with Parkinson’s disease impairs WASH complex association and inhibits autophagy. Nat Commun 5, 3828 (2014).

51. J. Follett, S. J. Norwood, N. A. Hamilton, M. Mohan, O. Kovtun, S. Tay, Y. Zhe, S. A. Wood, G. D. Mellick, P. A. Silburn, B. M. Collins, A. Bugarcic, R. D. Teasdale, The Vps35 D620N mutation linked to Parkinson’s disease disrupts the cargo sorting function of retromer. Traffic 15, 230–244 (2014).

52. O. Kovtun, N. Leneva, Y. S. Bykov, N. Ariotti, R. D. Teasdale, M. Schaffer, B. D. Engel, D. J. Owen, J. A. G. Briggs, B. M. Collins, Structure of the membrane-assembled retromer coat determined by cryo-electron tomography. Nature 561, 561–564 (2018).

53. T. Eguchi, T. Kuwahara, M. Sakurai, T. Komori, T. Fujimoto, G. Ito, S. I. Yoshimura, A. Harada, M. Fukuda, M. Koike, T. Iwatsubo, LRRK2 and its substrate Rab GTPases are sequentially targeted onto stressed lysosomes and maintain their homeostasis. Proc Natl Acad Sci U S A 115, E9115–E9124 (2018).

54. T. Kuwahara, K. Funakawa, T. Komori, M. Sakurai, G. Yoshii, T. Eguchi, M. Fukuda, T. Iwatsubo, Roles of lysosomotropic agents on LRRK2 activation and Rab10 phosphorylation. bioRxiv, 2020.2008.2025.267385 (2020).

55. A. Bhattacharya, R. Mukherjee, S. K. Kuncha, M. E. Brunstein, R. Rathore, S. Junek, C. Munch, I. Dikic, A lysosome membrane regeneration pathway depends on TBC1D15 and autophagic lysosomal reformation proteins. Nat Cell Biol, (2023).

56. K. Ito, M. Araki, Y. Katai, Y. Nishimura, S. Imotani, H. Inoue, G. Ito, T. Tomita, Pathogenic LRRK2 compromises the subcellular distribution of lysosomes in a Rab12-RILPL1-dependent manner. FASEB J 37, e22930 (2023).

57. J. L. Daly, C. M. Danson, P. A. Lewis, S. Riccardo, L. Di Filippo, D. Cacchiarelli, S. J. Cross, K. J. Heesom, A. Ballabio, J. R. Edgar, P. J. Cullen, Multiomic Approach Characterises the Neuroprotective Role of Retromer in Regulating Lysosomal Health. bioRxiv, 2022.2009.2013.507260 (2022).

58. A. Ungewickell, C. Hugge, M. Kisseleva, S. C. Chang, J. Zou, Y. Feng, E. E. Galyov, M. Wilson, P. W. Majerus, The identification and characterization of two phosphatidylinositol-4,5-bisphosphate 4-phosphatases. Proc Natl Acad Sci U S A 102, 18854–18859 (2005).

59. S. Takemasu, K. Nigorikawa, M. Yamada, G. Tsurumi, S. Kofuji, S. Takasuga, K. Hazeki, Phosphorylation of TMEM55B by Erk/MAPK regulates lysosomal positioning. J Biochem 166, 175–185 (2019).

60. Y. Hashimoto, M. Shirane, K. I. Nakayama, TMEM55B contributes to lysosomal homeostasis and amino acid-induced mTORC1 activation. Genes Cells 23, 418–434 (2018).

61. B. Ruprecht, J. Zecha, D. P. Zolg, B. Kuster, High pH Reversed-Phase Micro-Columns for Simple, Sensitive, and Efficient Fractionation of Proteome and (TMT labeled) Phosphoproteome Digests. Methods Mol Biol 1550, 83–98 (2017).

62. J. Cox, M. Mann, MaxQuant enables high peptide identification rates, individualized p.p.b.-range mass accuracies and proteome-wide protein quantification. Nat Biotechnol 26, 1367–1372 (2008).

63. S. Tyanova, T. Temu, P. Sinitcyn, A. Carlson, M. Y. Hein, T. Geiger, M. Mann, J. Cox, The Perseus computational platform for comprehensive analysis of (prote)omics data. Nat Methods 13, 731–740 (2016).

64. H. Ashkenazy, S. Abadi, E. Martz, O. Chay, I. Mayrose, T. Pupko, N. Ben-Tal, ConSurf 2016: an improved methodology to estimate and visualize evolutionary conservation in macromolecules. Nucleic Acids Res 44, W344–350 (2016).

65. X. Robert, P. Gouet, Deciphering key features in protein structures with the new ENDscript server. Nucleic Acids Res 42, W320–324 (2014).

